# Chromosome-level Genome Assembly of a Regenerable Maize Inbred Line A188

**DOI:** 10.1101/2020.09.09.289611

**Authors:** Guifang Lin, Cheng He, Jun Zheng, Dal-Hoe Koo, Ha Le, Huakun Zheng, Tej Man Tamang, Jinguang Lin, Yan Liu, Mingxia Zhao, Yangfan Hao, Frank McFraland, Bo Wang, Yang Qin, Haibao Tang, Donald R McCarty, Hairong Wei, Myeong-Je Cho, Sunghun Park, Heidi Kaeppler, Shawn M Kaeppler, Yunjun Liu, Nathan Springer, Patrick S Schnable, Guoying Wang, Frank F White, Sanzhen Liu

## Abstract

The highly embryogenic and transformable maize inbred line A188 is an attractive model for analyzing maize gene function. Here we constructed a chromosome-level genome assembly of A188 using long reads and optical maps. Genome comparison of A188 with the reference line B73 identified pervasive structural variation, including a 1.8 Mb duplication on the *Gametophyte factor1* locus for unilateral cross-incompatibility and six inversions of 0.7 Mb or greater. Increased copy number of the gene, *carotenoid cleavage dioxygenase 1* (*ccd1*) in A188 is associated with elevated expression during seed development. High *ccd1* expression together with low expression of *yellow endosperm 1* (*y1*) condition reduced carotenoid accumulation, which accounts for the white seed phenotype of A188 that contrasts with the yellow seed of B73 that has high expression of *y1* and low expression of the single-copy *ccd1*. Further, transcriptome and epigenome analyses with the A188 reference genome revealed enhanced expression of defense pathways and altered DNA methylation patterns of embryonic callus.

---

The maize inbred line A188 was derived from a line related to the commercial maize variety Silver King and a northwestern dent line ^1^. A188 is amenable to somatic embryogenic culture (**Supplementary Fig. 1**) and regeneration and was the first maize line used to produce genetically modified plants ^2^. A popular maize transformation line, Hi-II, was isolated from offspring of a cross between A188 and B73 ^3,4^. Although highly valuable for plant regeneration and transformation, A188 is not agronomic competitive, having small ears and low grain yield (**Fig. 1, Supplementary Table 1**). The line also exhibits a high degree of growth habit plasticity in response to varying environments. In particular, A188 is overly sensitive to abiotic and biotic stresses, including drought, heat, and bacterial and fungal diseases, in comparison to elite maize lines ^5^. Nonetheless, hybrids of A188 and B73 exhibit extensive heterosis (**Supplementary Fig. 2**). A188, therefore, in addition to traits related to transformability, can serve as a model inbred line for the genetic dissection of many important agronomic traits, heterosis, and plant-environment interactions.

**Fig. 1.**
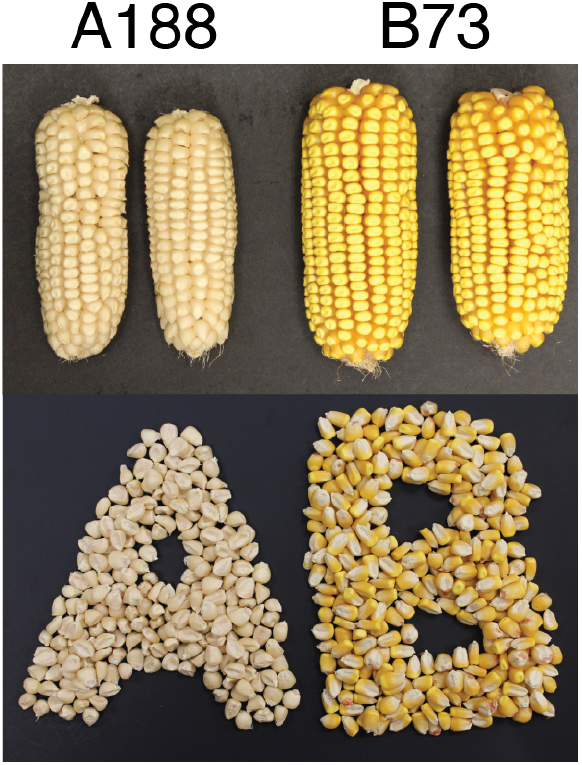
Seed photos of A188 and B73.

Efforts have been pursued to develop efficiency and quality strategies for maize genome sequencing and assemblies. The first maize reference genome for B73 was sequenced and assembled using bacterial artificial chromosomes (BACs) ^6^. Since then, additional assemblies have been produced using so-called next generation high throughput sequencing, including both short and long read technologies ^7–13^. Recently, two long-read technologies, PacBio and Nanopore, were combined with optical DNA mapping to produce a high-continuity maize assembly ^14^. Here, we used Nanopore long reads and optical DNA mapping to construct a chromosome-level maize genome of A188 for discovery of structural variation as well as performed transcriptomes and DNA methylome analyses of embryogenic callus.

## Chromosome-level A188 assembly

Long reads, representing a 90X coverage, were generated from A188 genomic DNA using the Oxford Nanopore sequencing platform. The N50 of read lengths is 23.9 kb, and the longest read is 264.5 kb (**Supplementary Fig. 3**). Genome assembly, performed using Canu, resulted in 1,830 contigs, comprising approximately 2.2 Gb of total sequences. The N50 of contigs is 5.99 Mb (**Supplementary Table 2**).

Read depths for contigs were assessed using Illumina short reads generated independently from seedling and immature ear DNAs to identify potential contamination from organelle genomes or extraneous microbial DNA whose contigs were expected to have differential read depths between the two tissues. Based on this strategy, contigs identified as the chloroplast or mitochondrion sequences were replaced respectively with the previously complete assemblies of A188 organelle genomes ^15,16^ and contigs from extraneous contamination were discarded (**Supplementary Fig. 4**). The remaining contigs were polished using raw Nanopore data and 80X PCR-free Illumina 2×250 paired-end whole genome sequencing reads (**Supplementary Table 3**), followed by the scaffolding with 113 A188 Bionano Genome (BNG) optical maps, for which the total length is 2.17 Gb and the N50 of these maps is 103.4 Mb. The BNG aided assembly placed 875 contigs into 39 scaffolds, which consist of 2.15 Gb. Chromosome pseudomolecules were then generated using a genetic map constructed from 100 B73xA188 double haploid (DH) lines. The final assembly (A188Ref1) consists of 2.25 Gb, including 10 chromosomal pseudomolecules, a mitochondrial genome, a chloroplast genome, and 986 scaffolds or contigs (**Table 1**).

**Table 1.**
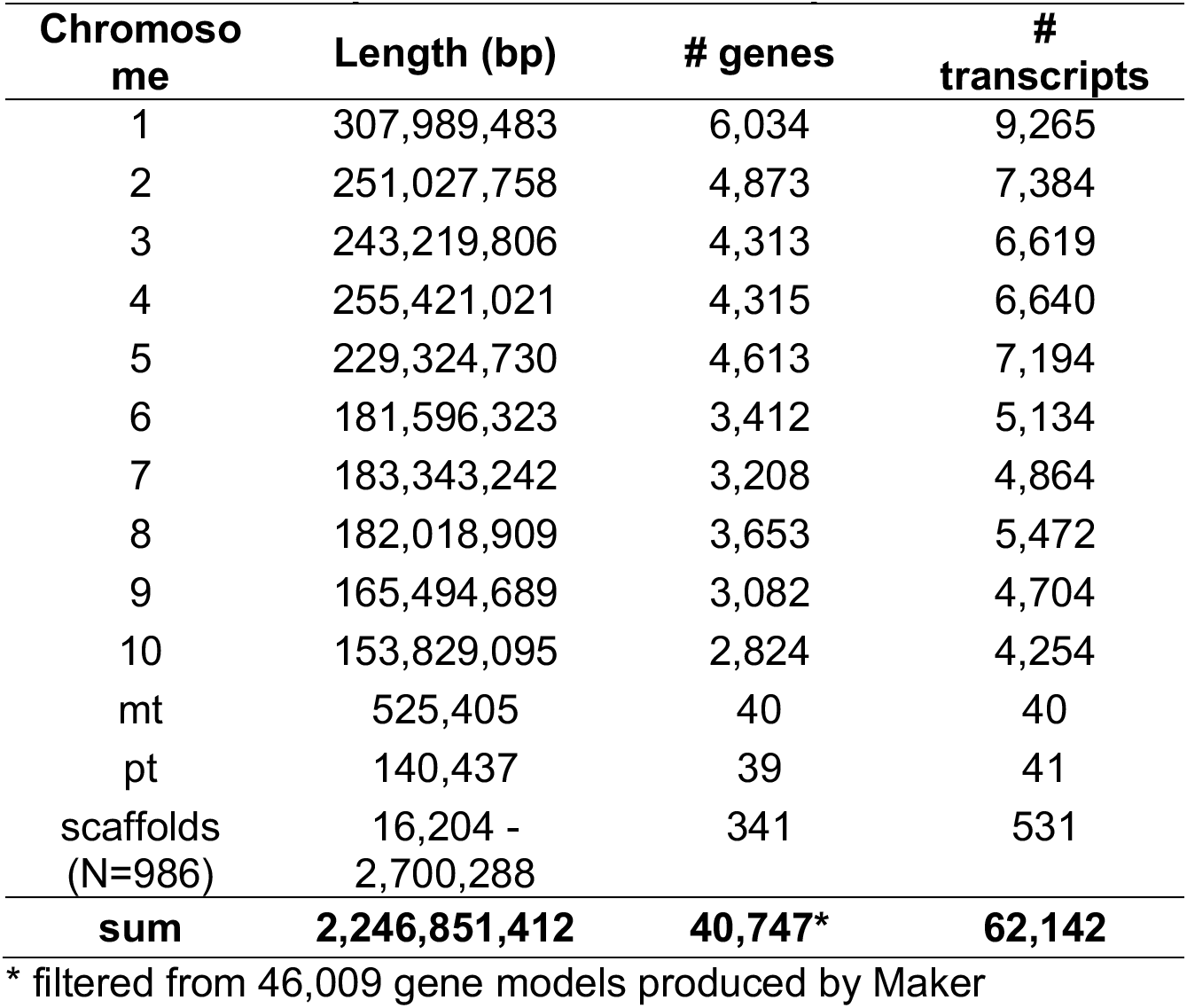
Summary of A188Ref1 assembly and annotation

| Chromosome | Length (bp) | # genes | # transcripts |
| --- | --- | --- | --- |
| 1 | 307,989,483 | 6,034 | 9,265 |
| 2 | 251,027,758 | 4,873 | 7,384 |
| 3 | 243,219,806 | 4,313 | 6,619 |
| 4 | 255,421,021 | 4,315 | 6,640 |
| 5 | 229,324,730 | 4,613 | 7,194 |
| 6 | 181,596,323 | 3,412 | 5,134 |
| 7 | 183,343,242 | 3,208 | 4,864 |
| 8 | 182,018,909 | 3,653 | 5,472 |
| 9 | 165,494,689 | 3,082 | 4,704 |
| 10 | 153,829,095 | 2,824 | 4,254 |
| mt | 525,405 | 40 | 40 |
| pt | 140,437 | 39 | 41 |
| scaffolds (N=986) | 16,204 - 2,700,288 | 341 | 531 |
| **sum** | **2,246,851,412** | **40,747\*** | **62,142** |

\* filtered from 46,009 gene models produced by Maker

The base accuracy of the A188Ref1 assembly was estimated at approximately 99.82% using the KAD pipeline ^17^. Approximately 96.4% of the potential errors are in transposons or other repetitive sequences. The estimated accuracy of genic sequences was >99.97%. The completeness of the A188 assembly was assessed using the BUSCO software ^18^ and found to contain 97.25% (3,189/3,278) of the Liliopsida core gene set, similar to the 97.36% (3,193/3,278) in the B73 reference genome (B73Ref4) ^7^.

## Presence of complex repeats and nuclear organelle sequences in A188Ref1

In total, 86.3% of the A188 genome sequence is annotated as repetitive elements. The long terminal repeat (LTR) retrotransposons *Gypsy* and *Copia* were the most prevalent elements, consisting of 44% and 23.9% of A188Ref1, respectively (**Fig. 2**). LTR centromere retrotransposon of maize (CRM) were largely co-localized with centromere-specific satellite repeat CentC, both of which were largely syntenic to the B73 centromeres ^7^. Approximately 8.3% of A188Ref1 is annotated as DNA transposable elements (TEs), including helitron and Miniature Inverted-repeat Transposable Elements (MITEs) (**Fig. 2**). Major knob clusters were found on the long arm of chromosomes 5 (5L), the short arm of chromosome 6 (6S), 7L, and 8L, and major subtelomeric repeats (4-12-1) were clustered on the distal regions of 1S, 3S, 4S, 5S, and 8L (**Fig. 2**). Through similarity alignments, we identified the 45S and 5S ribosomal DNA (rDNA) clusters on 6S and 2L, respectively (**Fig. 2**). Knob and rDNA locations were in agreement with previously reported A188 fluorescent in situ hybridization (FISH) data ^19^. Most repetitive components were located in regions of low-recombination contexts except the 5S rDNA locus and subtelomeric clusters (**Supplementary Fig. 5**).

**Figure 2.**
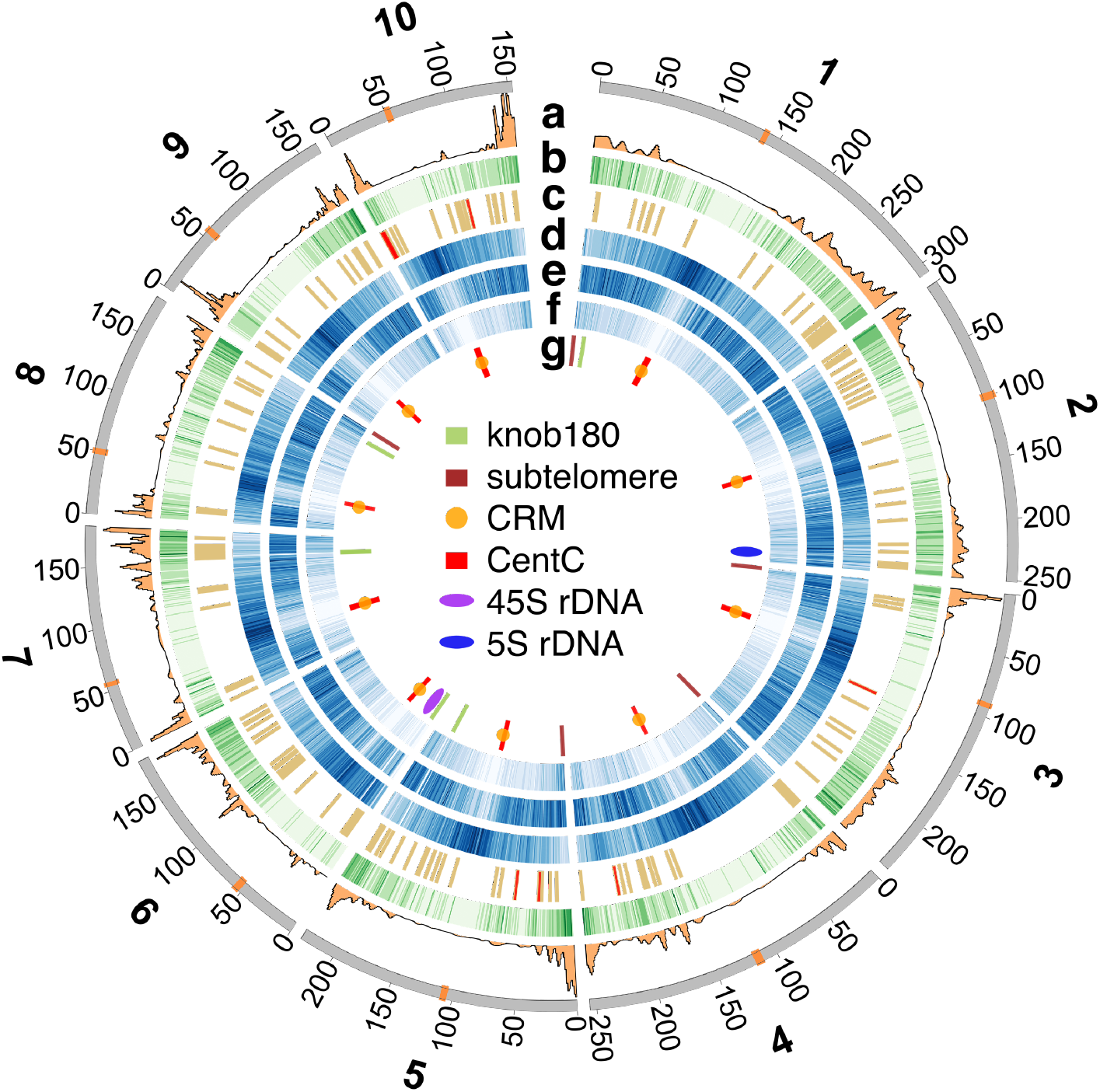
Circos plot of genomic features. Features on chromosomes 1 to 10 are: a) recombination rate (cM/Mb); b) gene density per Mb; c) gene clusters; d) number of *Gypsy* per Mb; e) number of *Copia* per Mb; f) number of MITEs per Mb; g) high-copy repetitive elements. The central inset is the legend for the track of g. Tracks of b, d, e, f are intensity-coded. The higher the intensity, the higher frequency each element. Centromeres are in orange on the outmost chromosome track, on which numbers are coordinates in Mb.

Nuclear mitochondrial DNA (NUMT) and nuclear plastid (chloroplast) DNA (NUPT) were identified at 10 and 21 genomic loci, respectively (**Fig. 3a, Supplementary Fig. 6**). The largest nuclear organelle-like sequence (~136 kb) is a NUMT locus on the short arm of chromosome 8, which contains an array of DNA transposons likely inserted subsequent to the NUMT integration. FISH analysis corroborated the chromosome 8 location and confirmed a homolog on the distal end of 10S (**Fig. 3b,3c**). In summary, the genomic locations of repetitive sequences and nuclear organelle sequences are largely consistent with previous findings by FISH ^20,21^, supporting the large-scale correctness of A188Ref1.

**Figure 3.**
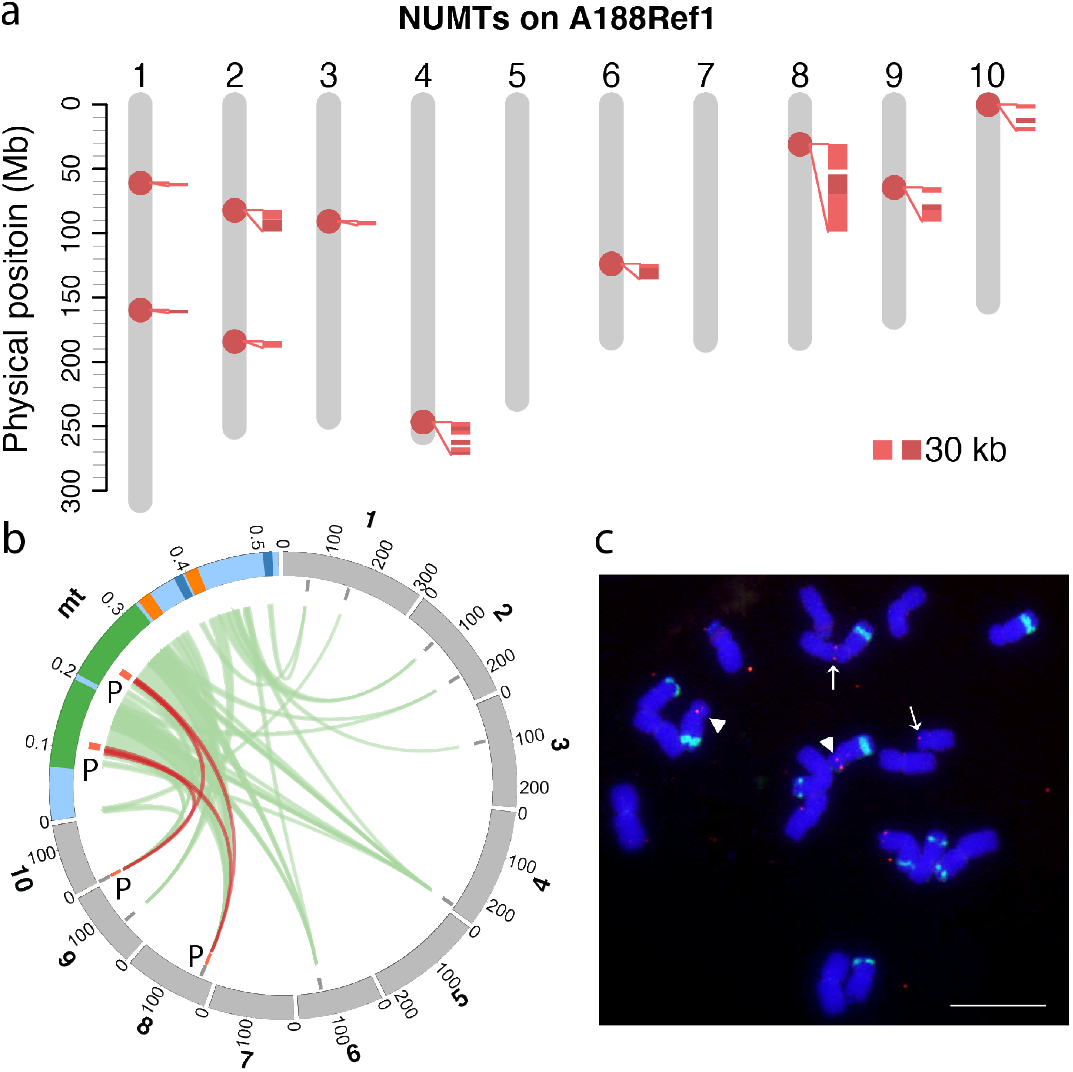
NUMT on A188 nuclear genomes. **a**). NUMT sequence on 10 chromosomes of A188Ref1. Each dot on chromosomes designates a potential NUMT integration. Close-up alignments with the mitochondrion (mt) genome are shown along NUMTs. Each alignment requires at least 5 kb match and 95% identity. **b**). Circos plot of alignments between the mt genome and ten chromosomes. The same color of green, orange, dark blue label duplicated regions in mt. “P” regions match the probe sequence used for FISH. Brown links highlight alignments on chromosomes 8 and 10. Note that the chromosomal scale is different from the mt scale. Numbers on the track are in Mb. **c**). Physical mapping of a mt DNA (mtDNA) and knob repeats on the mitotic metaphase chromosomes of maize A188. The knob repeat probe (green signals) was used to identify the chromosomes. Two FISH sites of mtDNA insertion on the chromosomes were detected: arrowheads, chromosome 8; arrows, chromosome 10. Bar=10 µm.

## Gene annotation

Annotation of A188Ref1 was performed using the Maker pipeline with evidence from transcripts assembled with A188 long Nanopore direct cDNA sequencing data, A188 RNA-Seq Illumina short reads, and transcripts from other maize lines, as well as protein sequences from closely related plant species. The Maker genome annotation resulted in 40,747 high-confidence gene models with 62,142 transcripts (A188Ref1a1) (**Table 1**). BUSCO evaluation showed that 97.8% Liliopsida conserved genes were in A188Ref1a1. Comparison of protein sequences identified 52,971 orthologous pairs between A188 and B73, consisting of 27,273 A188 genes and 27,529 B73 genes, as well as 18,521 pairs of paralogous genes in A188. We also identified 178 gene clusters each of which contains at least three paralogous genes, comprising in total 694 genes. The largest such cluster contains 25 genes putatively encoding pectin methylesterase (PME) on an unanchored scaffold. One cluster of 18 genes is located in the same unanchored scaffold sequence containing the aforementioned 25 PME genes; one cluster of nine PME genes is present on chromosome; and a cluster of five genes is on chromosome 5 (**Fig. 4a**). Gene clusters also include eight clusters of 42 nucleotide-binding leucine-rich repeat (NLR) disease resistance (R) genes (**Fig. 4a**). One NLR gene cluster on chromosome 10 has 16 genes homologous to the *rp1* gene that confers resistance to common rust ^22^ and was associated with Goss’s wilt resistance ^23^ (**Fig. 4b**). Most paralogous clusters were not located in regions with high recombination (**Fig. 4c**). Exceptions include the *rp1* locus, which has a high level of haplotype instability through frequent recombination among *rp1* paralogs ^24–26^. Divergent *rp1* haplotypes were observed between A188 and B73 that contains 11 *rp1* homologs at the syntenic locus (**Fig. 4b**).

**Figure 4.**
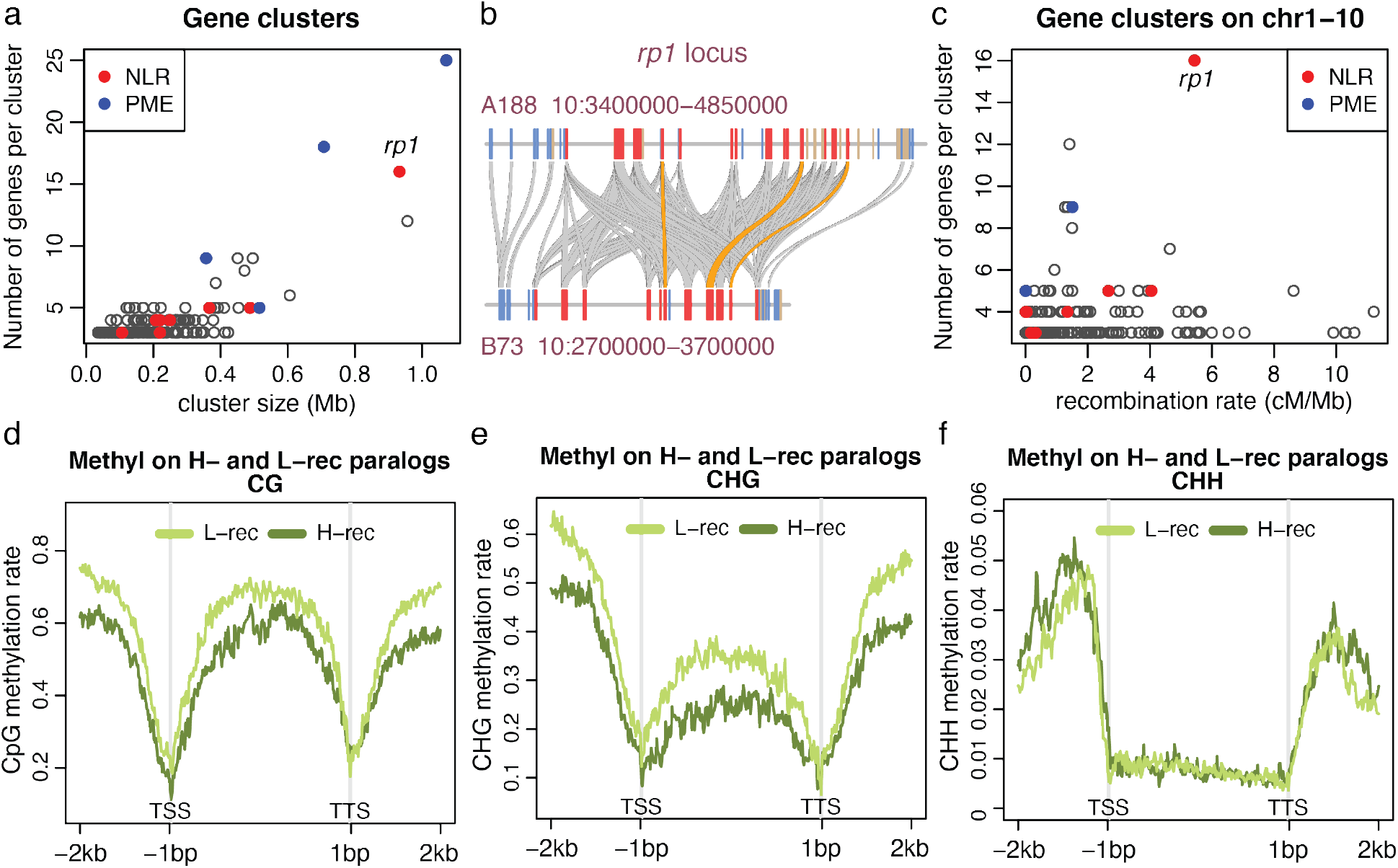
Gene clusters and paralogs in low- and high-recombination regions. **a**). The scatter plot of numbers of genes per cluster versus their cluster size. **b**). Example of an NLR gene (*rp1*) cluster in A188 and their alignments with the B73 *rp1* locus. Each rectangle box represents a gene with blue, tan, and red colors indicating plus, minus orientation, and *rp1* homologous genes. All *rp1* homologs are in the same minus orientation. Gray bands connect orthologs and orange bands highlight the top *rp1* alignments with at least 98.5% identity and a 2,500 bp match. **c**). The scatter plot of numbers of genes per cluster versus the recombination rate estimated 1 Mb around the midpoint of the cluster. All clusters plotted are on 10 chromosomes. **d-f**). Distribution of cytosine methylation in sequence contexts of CG, CHG, and CHH around paralogous genes. An average methylation rate per window across all examined genes from two replicates of seedling samples was determined and plotted versus the window order. A window in the gene body, from translation start site (TSS) to translation termination site (TSS), is 1/200 of the gene body in length. A window outside of the gene body is 20 bp.

We identified 2,259 paralogous gene pairs of which one member was located in a high-recombination chromosomal compartment and the other in a low-recombination compartment (Methods). Comparison of DNA methylation of A188 seedlings found that, on average, both CG and CHG methylation, where H represents A, C, or T, were higher near and within low-recombination paralogous genes as compared to high-recombination genes. No obvious differences were observed in CHH methylation (**Fig. 4d-f**). Comparison of gene expression between members of the paralogous pairs using seedling RNA-Seq data showed most paralogs had similar expression levels and no expression bias to either high-or low-recombination genes was observed for those paralogs that did exhibit differential expression (**Supplementary Fig. 7**). The result indicated that the genomic context of genes is a driver for a certain epigenomic modification but not a major driver for gene expression.

## High-level structural variation between A188 and B73

Structural variation between the A188 and B73 genomes was identified through comparisons of whole genome assemblies of both genomes using SyRI software (**Extended Data 1**) ^27^ and through the analysis of whole genome Illumina sequencing reads with CGRD (Comparative Genomic Read Depth) that is based on quantitative comparison of depths of short reads (**Extended Data 2, 3**) ^28^. SyRI revealed ~1.1 Gb of syntenic regions, 2,302 translocations, as well as 4,083 duplications in B73 and 2,333 duplications in A188 using a minimum cutoff of 10 kb for each translocation or duplication event (**Supplementary Table 4**). In addition, SyRI identified 441.9 Mb of B73 and 543.8 Mb of A188 DNA sequences that were not aligned with the other respective genome. Further filtering with CGRD that compared read depths between the two genomes, revealing 381.3 Mb of B73-specific sequences and 409.2 Mb of A188-specific sequences that represent presence and absence variance (PAV) or highly divergent sequences (HDS) (**Extended Data 4, 5**). These PAV/HDS regions contain 6,728 genes in B73 and 7,301 genes in A188. Gene ontology enrichment analysis indicated that genes related to endopeptidase inhibitor activity and extracellular activities are enriched in both PAV/HDS gene sets (**Supplementary Fig. 8**).

Seventeen large inversions of 0.5 Mb or greater were identified between the two genomes (**Fig. 5, Supplementary Fig. 9-17**, **Supplementary Table 5**). Nine of the seventeen inversions are likely errors in B73Ref4 as the newly released B73Ref5 (unpublished version 5 but available from maizeGDB) showed the same orientation as A188Ref1, including the largest inversion region (INV37083 on B73Ref4, 97.8-103.9 Mb on chromosome 4). FISH analysis of A188 and B73 corroborated the absence of inversion INV37083 (**Supplementary Fig. 18**). Recombination and pairwise linkage disequilibrium (LD, R^2^) values among single nucleotide polymorphisms (SNPs) within each inversion were determined, and out of eight remaining inversion candidates, six have recombination frequencies close to 0 and a high mean LD ranging from 0.56 to 0.79 of all pairs of SNPs that are separated by 0.2-0.3 Mb within an inversion, which are higher than the genome-wide average LD of 0.2 between SNPs in separated by 0.2 Mb (**Supplementary Table 5**). These six inversions exhibiting marked recombination suppression characteristic of inversion ^29^, therefore, are strongly supported. The six inversions range from 0.7 to 2.1 Mb in size, of which two are located close to the centromere of chromosome 2 and four are on 3L, 4L, 5L, and 9L (**Supplementary Table 5**). In total, the six inversion sequences harbor 69 B73 genes and 75 A188 genes. The syntenic relationships of these genes were largely maintained between inverted sequences in the two genomes (example in **Fig. 5d**), although the gene sequences are divergent in a high degree from each other (**Fig. 5b**, **Supplementary Fig. 10,11,12,16**). The divergence of these inversions indicated that the inversions were not recent events maintained in modern maize populations. Admixture structure analysis showed that both A188 and B73 haplotypes of 3/6 inversions exist in teosinte, the maize wild ancestor (**Supplementary Fig. 19**), and no clear evidence of the haplotype of the remaining three inversions exist in teosinte (**Supplementary Fig. 20**). Among landraces, both the A188 and B73 haplotypes of all six inversions could be identified.

**Figure 5.**
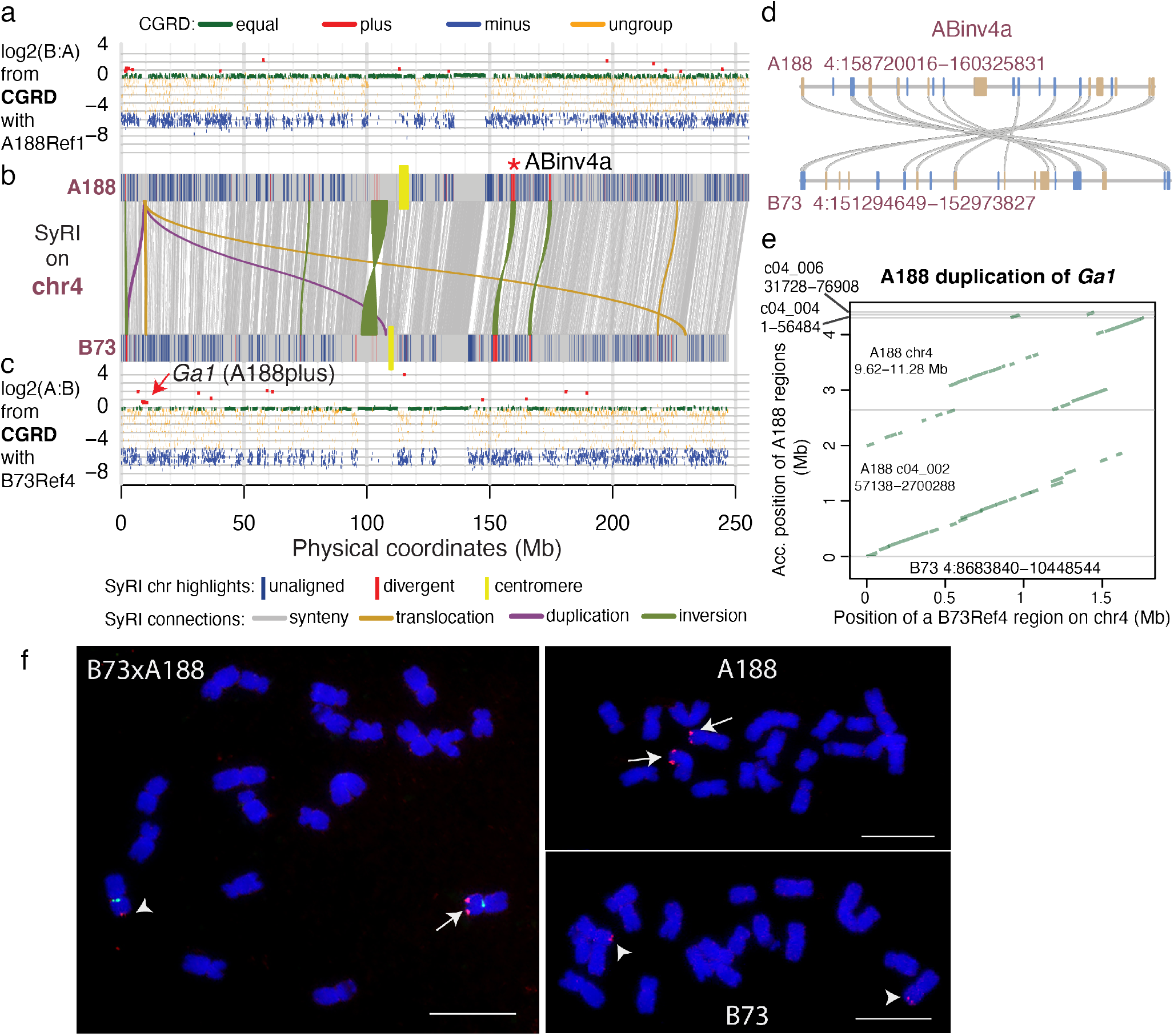
Megabase-level duplication and inversion on chromosome 4. a, b, c). SyRI and CGRD results on chromosome 4. a). The CGRD result using A188Ref1 as the reference genome. Y-axis represents log2 values of ratios of read depths of B73 to A188, log2(B:A), signifying copy number variation (CNV). Regions with higher and lower sequence depths of B73 versus A188 were B73 plus (red) and B73 minus (blue), respectively. Green and orange represents conserved and ungrouped regions, respectively. b). The SyRI result is displayed. Alignments of syntenic blocks larger than 10 kb and alignments of other rearrangements larger than 0.5 Mb are plotted. On each A188 and B73 chromosome, segments that were not aligned to the other genome or highly divergent with the other genome are highlighted. The red * labels a well-evidenced inversion. c) The CGRD result using B73Ref4 as the reference genome. The similar color scheme to that in a) is used. d). Synteny of genes (rectangle blocks) in the well-evidenced inversion (ABinv4a) regions between A188 and B73. blue and tan colors stand for plus and minus gene orientations. e). A dot plot between the 1.8Mb B73 region that was duplicated in A188 and its aligned regions in A188Ref1. f). FISH of the PME probe on A188, B73, and F_1_ (B73xA188). Cent4 probe (green) that specifically targeted on chromosome 4 centromere was used in F_1_ FISH. Arrows and arrow heads point at PME signals of A188 and B73 chromosomes, respectively. Bar=10 μm.

CGRD analysis also identified a A188 duplication of a 1.8 Mb region from 8.68 Mb to 10.45 Mb on chromosome 4 of B73Ref4 (**Fig. 5c**). In A188, a portion of the duplication was found in the unanchored scaffold c04_002 while most of the remaining duplicated sequences can be found in chromosome 4 (**Fig. 5e**). The duplication region overlapped with the *Gametophyte factor1* (*Ga1*) locus conferring unilateral cross-incompatibility (UCI) ^30^. The underlying causal gene of B73, Zm00001d048936, encodes a PME, which is a wildtype allele. We designed a PME DNA probe that is from the duplication and repeatedly matches 35 loci in B73Ref4 and 78 loci in both the region on chromosome 4 and the scaffold c04_002 on A188Ref1. FISH using this probe resulted in strong hybridization signals on A188 4S and weak signals on B73 4S, indicating that the duplication occurred locally on 4S (**Fig. 5f**). The B73 Zm00001d048936 gene has no additional homologous copies in B73Ref4 but five homologous sequences can be identified on the duplicated sequence of A188Ref1, including the syntenic gene Zm00056a022745 that is identical to Zm00001d048936. Collectively, the result documented the complexity and the potential dynamic of the *Ga1* locus of maize.

## Associating structural variation with phenotypic variation

The CGRD result indicated that A188 had many more copies (A188plus) at a region from 155.23 to 155.24 Mb of chromosome 9 in B73Ref4 (**Supplementary Fig. 15**). This region includes the *carotenoid cleavage dioxygenase 1* (*ccd1*) gene catalyzing the cleavage of carotenoids to apocarotenoid products, which is located at the White cap locus (*Wc1*) conditioning kernel colors ^31^. SyRI analysis supported a duplication of this region but failed to find a number of copies in A188. SyRI analysis also indicated the duplicated region is embedded in A188-specific sequences (**Fig. 6a**). Comparison of A188Ref1 with an A188 BNG optical map aligned to the duplication region indicated the incomplete assembly of the region. Previously, tandem repeats of an ~27 kb sequence at the *Wc1* locus were discovered ^32^. Each repeat exhibits four discernible sites that can be detected *via* Bionano analysis, referred to as Type A repeat. Analysis of A188 sequences revealed a repeat variant containing an additional site, referred to as Type B repeat. Based on the BNG map, the A188 genome contains 13 intact tandem copies of the 27 kb sequence, consisting of 9 copies of Type A and 4 copies of Type B repeats, as well as partial copies of the 27 kb sequence on both ends of the array. Each repeat copy contains a *ccd1* gene, indicating at least 13 copies of *ccd1* in A188 (**Fig. 6b**), consistent with the A188plus result from the CGRD analysis. Neither intact Type A nor B repeat exists in B73, which, however, does contain a *ccd1* gene.

**Figure 6.**
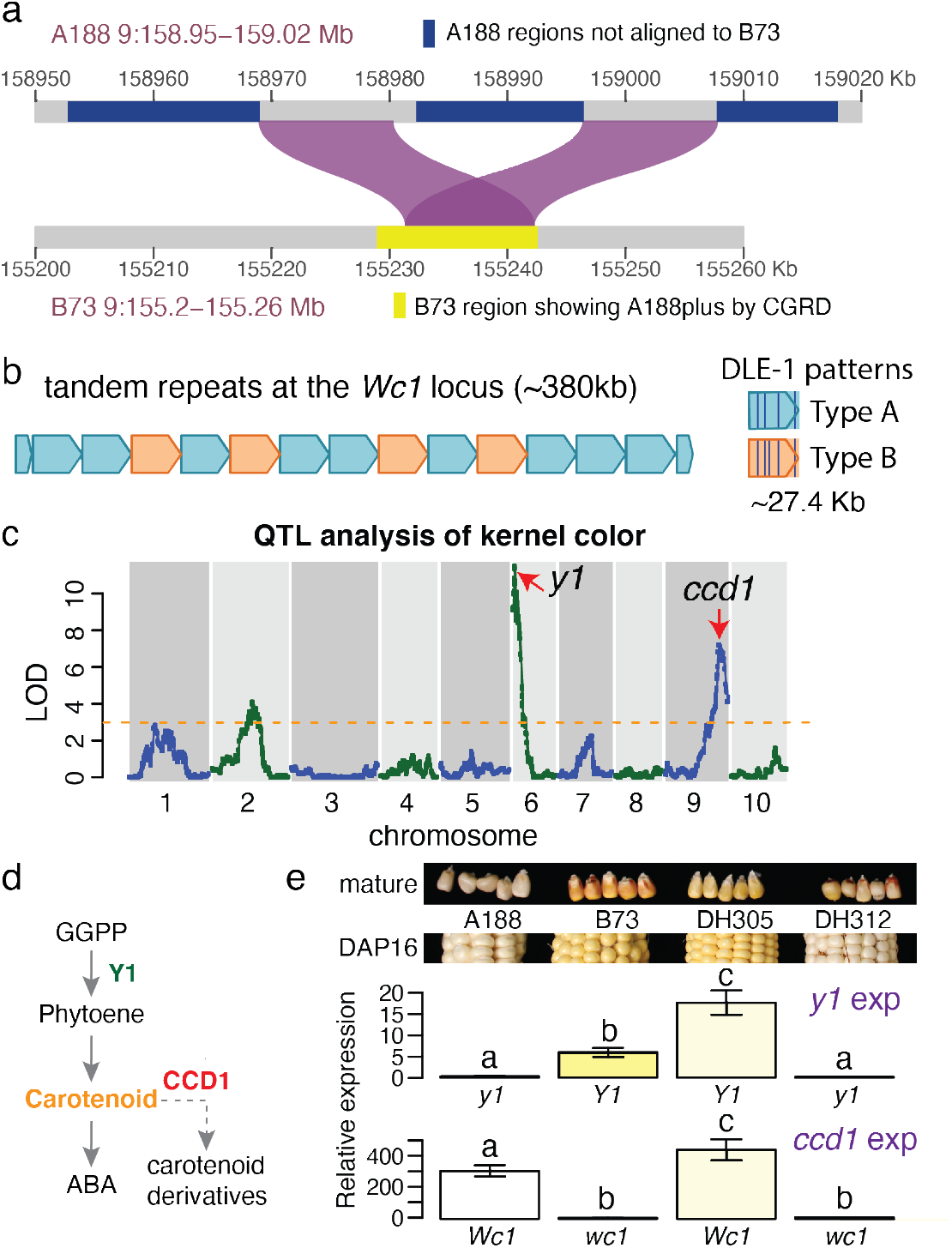
Structural variation and genetic analysis of the *Wc1* locus. a). Duplication alignments between A188 and a B73 region identified as A188plus by CGRD. b). Tandem repeats of 13 intact copies of 27.4 kb sequences. Two DLE-1 restriction patterns in repeat units: Types A and B were identified. c). The QTL result of kernel color using the DH population. Arrows point at locations of known causal genes. d). A simplified carotenoid pathway. GGPP stands for geranylgeranyl diphosphate. e) Seeds at 16 days after pollination (DAP16) were collected and used for quantifying gene expression (exp) of *y1* and *ccd1*. Three biological replicates were used. Bars are color-coded based on colors of mature seeds. Error bars represent standard variation. Letters on top of bars are statistical groups determined by Tukey tests. *Y1* (*wc1*) and y1 (*Wc1*) stand for B73 and A188 alleles, respectively. Mature seeds from the same lines show slightly different colors from seeds of DAP16.

A188 seeds are white, whereas B73 seeds are yellow (see **Fig. 1**). Analysis of quantitative trait locus (QTL) of kernel colors of B73xA188 DH lines resulted in two major QTLs on chromosomes 6 and 9, both of which were discovered in a previous genome-wide association study ^33^, as well as a weaker peak on chromosome 2 (**Fig. 6c, Supplementary Fig. 21**). Two known genes *y1* and *ccd1* in the major peaks are responsible for kernel colors (**Fig. 6d**) ^32,34^. The dominant *Y1* allele conditions yellow kernels ^34^. Several variants exist between the A188 *y1* (Zm00056a032392) and B73 *Y1* (Zm00001d036345) alleles, including one amino acid polymorphism (Ser258Thr) in the coding region (**Supplementary Fig. 22**) and polymorphisms found in 5’ upstream and 3’ downstream regions, including a (CCA)_n_ microsatellite variation in the 5’ untranslated region ^35^ (**Supplementary Fig. 23**). Quantitative reverse transcription PCR (qRT-PCR) reveals higher expression of the *y1* gene in B73 relative to A188 (**Fig. 6e**). In contrast, the B73 *ccd1* expression was much lower than that of A188, presumably due to the differences in copy number (**Fig. 6e**). Because higher expression of functional alleles of the *ccd1* and *y1* genes is expected to reduce and increase the accumulation of carotenoids, respectively, the differences in the expression of the *ccd1* and *y1* genes in B73 and A188 likely explains yellow kernels of B73 and white kernels of A188 (**Fig. 6d,6e**). The expression levels of the alleles in two DH lines with different allele combinations of these two loci were similar (**Fig. 6e**).

## Distinct gene expression and hypermethylation in calli relative to seedlings

Transcriptomic data were generated for 11 diverse tissues with three biological replicates each. Both principal component analysis and clustering of these tissue samples based on their genome-wide gene expression showed that the callus from tissue culture were closely related to root, leaf base, embryo, and ear, but distinct from middle leaf, leaf tip, and seedlings (**Supplementary Fig. 24, Extended Data 6**). A set of 734 callus featured genes were identified that exhibited at least 2-fold up-regulation in the callus as compared to any other tissues (**Supplementary Table 6**). Genes involved in cell wall biosynthesis, defense activity, heme binding, transmembrane transport, and transcription regulation are enriched in these featured genes (**Supplementary Fig. 24**). The top six enriched transcription factor families are WOX, AUX/IAA, LBD, AP2, WRKY, and NAC, which included homologs of *Baby boom* (AP2) and *Wuschel2* (Wox) genes relevant to cell division and expansion (**Supplementary Fig. 25**) ^36^.

The callus and seedling tissues were selected for examination of genome-wide DNA methylation levels. The callus exhibited elevated methylation for all three sequence contexts as compared to the seedling, 89.3% vs 85.2% on CG, 74.5% vs 71.9% on CHG, and 3.2% vs 1.5% on CHH (**Supplementary Table 7**). The analysis of CG and CHG methylation over all genes did not find major differences between callus and seedling tissue (**Fig. 7a, 7b**). However, there were major differences in the level of CHH methylation (**Fig. 7c**). While the shape of the profile is similar in seedling and callus tissue the level of CHH methylation is much higher within and surrounding genes (**Fig. 7c**). On average, there were no major changes in the level of CG or CHG methylation over repetitive elements but there was a consistent trend for slightly higher CG methylation callus for most classes of repetitive elements (**Fig. 7d, 7e**). Similarly, CHH methylation was slightly higher for most classes of repetitive elements with the most notable increase observed at MITE elements (**Fig. 7f**).

**Figure 7.**
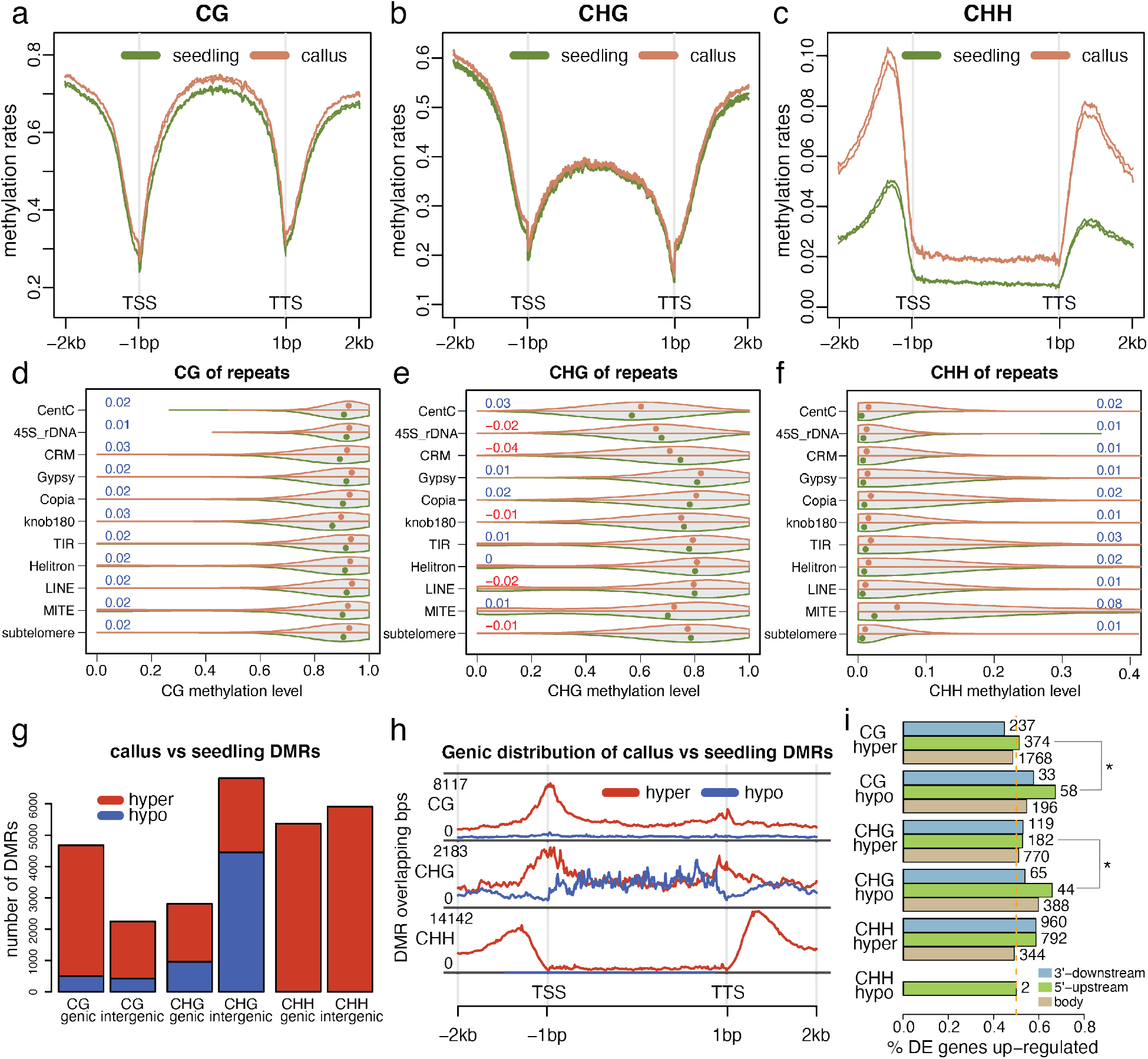
DNA methylation in callus and seedling tissues. (**a, b, c**). Distribution of cytosine methylation in three sequence contexts (CG, CHG, and CHH) around genes in two biological replicates of the callus (orange) and two biological replicates of the seedling (green). An average methylation rate per window across all examined genes was determined and plotted versus the window order. A window in the gene body is 1/200 of the gene body. A window outside of the gene body is 20 bp. (**d, e, f**). Violin plots of methylation on repetitive sequences. For each violin plot, the top half is the distribution of methylation in the callus and the bottom half is the distribution of methylation in the seedling. Each dot represents the median of methylation rates. Numbers stand for the mean methylation differences between the callus and the seedling, which are color coded with blue and red to represent increased methylation and decreased methylation in the callus, respectively. All differences are significant (p-value<0.0001) by paired t-test. (**g**) Barplots of DMRs on genic regions, including 2 kb beyond each of TSS and TTS, and the rest of the genome (intergenic regions). (**h**) Distribution of DMR sequences around genes. The definition of the gene body is the same as described in **a**-**c**. (**i**) Proportions of DE genes up-regulated in hyper DMR and hypo DMR regions (gene body, 1 kb 5’ upstream and 1 kb 3’ downstream regions). Numbers on top of bars are numbers of DE genes. Stars indicated significances (p<0.05) from χ# tests for the independence of the DMR and DE changing directions. In **g**, **h**, and **i,**hyper and hypo stand for increased and decreased methylation in the callus relative to the seedling, respectively.

Differentially methylated regions (DMRs) were identified through comparison of the DNA methylation profiles of callus and seedling. In total, 6,927 CG DMRs, 9,631 CHG DMRs, and 11,275 CHH DMRs were identified with the mean lengths of 200 bp, 243 bp, and 272 bp, respectively (**Supplementary Table 8**, **Extended Data 7, 8, 9**). Hypermethylation in callus relative to seedling was the predominant type of DMRs for both CG and CHH methylation in both genic and intergenic regions while CHG exhibited roughly equal proportions of hyper and hypomethylation DMRs with more hypermethylation in genic regions and more hypomethylation in intergenic regions (**Fig. 7g**). The analysis of the distribution of DMRs relative to genes revealed that the CG DMRs were enriched near TSS regions while CHH DMRs tended to be found in regions just upstream or downstream of genes, mirroring CHH island distributions (**Fig. 7h**) ^37,38^. CHG DMRs exhibited different trends for localization for hyper and hypomethylated DMRs with hypermethylation DNAs enriched at TSS and TTS regions and hypomethylated DMRs enriched in gene bodies (**Fig. 7h**). The high frequency of some types of DMRs near the TSS led us to assess whether these DMRs may be contributing to differential expression in callus relative to seedling tissue. Genes with DMRs were enriched for being differential expression (DE) in seedling relative to callus compared to genes without DMRs (*χ*^2^=20.9, p-value=4.9e-6). Based on prior studies in maize we expected that gains of CG or CHG methylation near the TSS would be associated with down-regulation of expression while gains of CHH upstream of the promoter might be associated with up-regulation of expression ^39,40^. We found that the DE genes with hypomethylated or hypermethylated DMRs at most regions exhibited roughly similar numbers of up-and down-regulated with exception at CG hypomethylation at 5’ upstream regions of genes and CHG hypomethylation in the gene body, both of which were associated with up-regulation of gene expression in the callus (**Fig. 7i, Supplementary Table 9**). These results reveal dynamic changes in some types of DNA methylation in callus relative to seedling and a marginal association of DNA methylation with gene expression changes.

## DISCUSSION

A chromosome-level A188 genome was generated using Nanopore long reads, Illumina short reads, and Bionano optical maps, which added a new reference genome to the collection of sequenced maize genomes ^6–14^. The strategy of comparing read depths of Illumina data from independent DNA sources to filter contigs before the scaffolding eliminated the contamination of DNA sequences from organelle genomes or microorganisms, while preserving nuclear integrated organellar sequences. The strategy can, and probably should, be applied for other genome assembly projects to differentiate nuclear DNA sequences from any potential contaminants that are subject to change in contamination levels in samples from different sources.

We also employed an approach, CGRD, for the discovery of genome structural variation based on quantitative comparison of depths of sequencing reads. Detailed characterization of genomic structural variation in complex genomes such as maize is challenging. Comparison using complete genome sequences based on their alignments would be an ideal method to reveal copy number variation and rearrangements. However, technically, alignment-based methods still need to overcome the complexity due to repetitive sequences. More critically, finding structural variation with assembled sequences is subject to the quality of assemblies. Unfortunately, assemblies of most plant genomes or other large complex genomes are generally not complete or error-free. For example, B73Ref4 appears to miss the topmost region of the short arm sequence of chromosome 6 (**Supplementary Fig. 13**) and includes multiple assembly inversion errors. CGRD based on comparison of depths of short reads complements the approaches that rely on whole genome comparisons such as SyRI ^27^. In particular, the CGRD pipeline can detect copy number variation missed by SyRI due to incomplete assembly at structurally complex regions. For example, we identified a 1.8 Mb A188 duplication at the *Ga1* locus and a high-copy tandem duplication of *Wc1* in A188 through CGRD, both of which did not stand out from SyRI analysis. The two methods are complementary in that CGRD captures unbalanced structural variation such as copy number variation rather than balanced structural variation such as translocations or inversions, which SyRI can detect. Therefore, the combination of SyRI and CGRD provides an optimal strategy for discovery of genomic structural variation, which is critical for further characterizing their impacts on gene expression and phenotypes.

In addition to the detection of large duplications and inversions, analysis of structural variation elucidated a tandem structure surrounding the *ccd1* gene that consists of 13 repeats in the A188 genome. Consistent with the previous findings, high copy number of *ccd1* corresponds to high expression, which presumably leads to a high activity of the carotenoid cleavage enzyme, resulting in enhanced carotenoid degradation ^32^. The expression of the A188 *y1* allele during seed development is low, while the *y1* expression in B73 is relatively high ^41^. Both alleles were highly expressed in some non-seed tissues such as leaf. Although the A188 Y1 protein sequence contains a single amino acid polymorphism as compared to the B73 Y1 protein, the A188 *y1* allele is likely functional, producing a low but perceptible level of carotenoid at certain stages of seed development, as evidenced by pale yellow seeds of recombinant DHs that carry the *y1* allele of A188 and the single copy *ccd1* allele of B73. We have identified an additional minor QTL that was concordant with QTLs from multiple other B73-derived bi-parental populations ^42^. The candidate gene *zep1* (Zm00001d003513) encoding zeaxanthin epoxidase was previously identified but functional validation is needed ^43^. Based on kernel colors of DH lines, the three QTL loci are not sufficient to fully determine kernel colors. Analysis with a larger B73xA188 derived population could possibly find additional loci influencing kernel colors.

Plant tissue culture from a highly differentiated tissue to the callus is the process of dedifferentiation to gain pluripotency for at least some cells ^44^. For cells under tissue culture, the transition of differentiation status is a highly stressful procedure ^45^. Our transcriptomic data revealed that defense genes were enriched among the callus featured genes that were up-regulated in the callus as compared to any other tissues examined. A number of NLR and defense-related genes, including *Pathogenesis-related protein 1* (*PR1*) (Zm00056a001451), were activated in the callus. A high-level stress is considered to produce somaclonal variation that is, by definition, phenotypic variation largely due to genomic alterations induced by tissue culture ^46^. Our transcriptomic data indicated that the plant defense was activated during the callus formation from immature embryos, by which the genome protection might be reinforced to reduce DNA damage or genome rearrangements that cause somaclonal variation. Hypermethylation is considered to be a protection mechanism against stresses, which enhance genome stability and safeguard genome integrity ^47^. Our comparison between the callus and the seedling uncovered globally elevated methylation in the callus in all three sequence contexts. Consistently, hypermethylation in the callus relative to the immature embryo was found in a study that used another maize inbred line and methylated DNA immunoprecipitation sequencing (MeDIP-seq) data ^48^. In this study, 24 nt small RNA was shown to be positively correlated with DNA methylation. In rice, CG hypermethylation was seen in one- and three-year callus relative to the shoot in the mutant of rice *MET1-2* that is a major DNA methyltransferase in maintaining CG methylation, but in the wildtype, only CHH hypermethylation was observed ^45^. In our study, the callus versus seedling comparison showed that A188 *MET1-2* homolog Zm00056a035610 was ~2x up-regulated in the callus, and *mop1* (Zm00056a013519), a homolog of RNA-dependent RNA polymerase 2 that involved in the production of 24 nt small RNA ^49^, was 5-6x up-regulated in the callus, indicating that the transcriptomic machinery was regulated to enhance globe DNA methylation in the callus. In plants regenerated from calli, CG and CHG methylation tended to be lost as compared to non-regenerated plants and many were heritable ^50^. Similarly, heritable hypomethylation was revealed in regenerated plants in an earlier maize study ^51^. In rice, as compared to non-regenerated plants in rice, pronounced hypomethylation was found in regenerated plants from tissue culture ^52^. The discrepant DNA methylation levels between regenerated plants and calli indicated that most methylation gained from tissue culture is not stable or heritable. Collectively, DNA methylation was elevated during the formation of the callus, likely enhancing the cellular defense for the maintenance of genome integrity, but the majority of DNA methylation gained might be demethylated during re-differentiation, resulting in hypomethylated regenerated plants.

DH lines have been generated from A188 and B73, a transformation recalcitrant line. Genome-wide genotyping information of DH lines enables the projection of A188 and B73 genome sequences in each DH line. Transformable DH lines with selectable desired genetic background and the availability of A188 genome sequences, together with the B73 reference genome, will provide foundational community resources for genome engineering and functional genomic studies in maize.

## METHODS

### Genetic materials

A188 (PI 693339) seeds were obtained from the North Central Regional Plant Introduction Station in Ames, IA. The A188 inbred line was derived from a cross between the inbred line 4-29 and the inbred line 64, also named A48, followed by four generations of backcross with 4-294. The line 4-29 was derived from the commercial variety Silver King and the line 64 was from a northwestern dent line ^1^. Double haploid lines were developed from the F_1_ of B73 (PI 550473) x A188 at the Doubled Haploid Facility at Iowa State University.

### Nanopore A188 whole-genome sequencing

A188 were grown in the greenhouse at 28°C, with a photoperiod of 14:10h (light:dark). Nuclei were isolated from seedling leaves using a modified nucleus isolation protocol ^53^ and dissolved in buffer G2 (Qiagen). The lysate was used for DNA isolation with Qiagen DNeasy Plant Mini Kit (Qiagen) following the manufacturer protocol. A188 genomic DNA was size selected for 15-30 kb and above with the BluePippin cassette kit BLF7510 (Sage Science) high-pass-filtering protocol, followed by a library preparation with the SQK-LSK109 kit (Oxford Nanopore). Each DNA library was loaded on a FLO-MIN106D flowcell and sequenced on MinION (Oxford Nanopore). The basecaller Guppy (version 3.4.4) converted FAST5 raw data to FASTQ data with default parameters.

### Illumina A188 whole-genome sequencing

Three independent A188 leaf samples were collected for extracting nuclear DNAs. Two were used for PCR-free paired-end 2×125 bp Illumina sequencing and one was used for PCR-free paired-end 2×250 bp Illumina sequencing on Hiseq2500 at Novogene. In addition, genomic DNA was extracted from A188 immature ears for additional PCR-free paired-end 2×250 bp Illumina sequencing. Therefore, comparable 2×250 bp data were generated from the leaf and ear tissue samples. The 2×125 bp Illumina sequencing data were comparable with the previously generated 2×125 bp B73 whole genome sequencing data (SRR4039069 and SRR4039070) ^54^, both of which were used for CGRD analysis.

### Assembly of Nanopore data *via* Canu

FASTQ Nanopore data were assembled with Canu (1.9) with the following options: "'corMhapOptions=--threshold 0.8 --ordered-sketch-size 1000 --ordered-kmer-size 14' correctedErrorRate=0.105 genomeSize=2.4g minReadLength=10000 minOverlapLength=800 corOutCoverage=60".

### Contigs filtering

Leaf and ear 2×250 bp data were aligned to the contigs with the “mem” module in bwa (0.7.12-r1039) ^55^. Uniquely mapped reads with less than 15% mismatches were used to determine read count per contig with the “intersect” module of BEDTools (v2.29.2) ^56^. The log2 of the ratio of read counts normalized by using total reads of leaf and ear samples was calculated for each contig. The contigs with a log2 value larger than 0.5 were considered to the contigs with variable counts from leaf and ear samples. The contigs (N=21) that had variable counts and less than 100 kb and were not anchored to B73Ref4 *via* Ragoo (version 1.2) ^57^ were discarded. In addition, the contigs (N=16) smaller than 15 kb were also discarded.

Through analysis of read counts, the contigs that had variable counts and matched with the previously sequenced mitochondrion genome sequence (Genbank accession: DQ490952.1) and the chloroplast genome sequence (Genbank accession: KF241980.1) were identified. One chloroplast contig and 13 mitochondrion contigs were found. The chloroplast contig had almost identical sequences to KF241980.1. The failure of assembling mitochondrial contigs into one was likely due to heterogeneous forms of mitochondria. In A188Ref1, the previously assembled, DQ490952.1 and KF241980.1, were used to represent the mitochondrion and chroloplast genomes, respectively.

### Sequence polishing of assembled contigs

After filtering contigs that were derived from organelles or contamination, the remaining contigs were first polished with raw Nanopore reads that contained signal information using Nanopolish (0.11.0) (github.com/jts/nanopolish). Briefly, Nanopore reads were aligned with the contigs using the aligner Minimap2 (2.14-r892) ^58^. Polymorphisms, including small insertions and deletions as well as single nucleotide polymorphisms, were called and corrected. The Nanopolish polishing was performed twice, followed by twice polishing with Illumina sequencing data using Pilon (version 1.23) ^59^. In each Pilon polishing, reads were aligned to contigs with the module of “mem” in bwa (0.7.12-r1039) ^55^. Contigs were corrected with the parameters of (--minmq 40 --minqual 15) using Pilon.

### Hybrid scaffolding with Bionano data and polished contigs

Bionano raw molecules were filtered for removing molecules less than 100 kb. The remaining molecules were assembled into Bionano maps with the assembly module in the software Bionano Tools (v1.0). Five times extension and merge iterations and noise parameters were automatically determined by using the parameters of “-i 5 -y”. The hybrid scaffolding module from the Bionano Tools was used for scaffolding polished contigs. The conflict filter level for both genome maps and sequences were set to 2 by using the parameters of “-B 2 -N 2”.

### Construction of a B73xA188 genetic map

Genomic DNA of DH lines was extracted by using BioSprint 96 DNA Plant Kit (Qiagen), and normalized to 10 ng/uL for Genotyping-By-Sequencing (GBS) modified from tunable GBS ^60^. Briefly, for each genomic DNA, the restriction enzyme *Bsp1286I* (NEB) was used for DNA digestion for 3 hrs at 37^°^C, followed by ligation with a barcoded single-stranded oligo with T4 DNA ligase (NEB) for 1 hrs at 16^°^C. Enzymatic activity was inactivated at 65^°^C for 20 min and all samples of ligated DNA were pooled, followed by purification with Qiagen PCR purification kit (Qiagen). The purified ligated DNA was subject to PCR amplification with Q5 high-fidelity DNA polymerase (NEB), followed by purification with Agencourt AMPure XP (Beckman Coulter). The final sequencing library product was prepared by size selection at the range of 200 to 400 bp by a Pippin Prep run with 2% agarose gel cassettes (Sage Science). Illumina sequencing was performed on a HiseqX 10 at Novogene.

Raw FASTQ data were deconvoluted to multiple samples and trimmed to remove barcode sequences and low-quality bases with Trimmomatic (version 0.38). Clean reads were aligned to polished contigs with the “mem” module of bwa and uniquely mapped reads with less than 8% mismatches were used for SNP analysis. SNPs were discovered by HaplotypeCaller of GATK (version 4.1.0.0) and filtered by SelectVariants of GATK to select biallelic variants ^61^. SNP sites with at most 80% missing data, at least 10% minor allele frequency and at most 5% heterozygous rates remained. A segmentation (or binning) algorithm was implemented to determine genotypes of chromosomal segments in each DH line ^60^. Genotypes of bin markers of 100 DH lines were used to construct a genetic map with MSTmap ^62^.

Another genetic map (**Extended Data 10**) was built using A188Ref1 as the reference genome with 37 additional DH lines. Recombination data was inferred from the genetic map (BAgm.v02) based on A188Ref1.

### ALLMaps to build pseudomolecules

The genetic map that was built based on polished contigs as the reference genome was used for further scaffolding. Each scaffold harbored more than 10 markers. In total, 29 scaffolds were on the map. Scaffolds were aligned to B73Ref4 *via* NUCMer. Based on the orientation of scaffolds relative to B73Ref4 chromosomes, the order of markers in each linkage group was either kept the same order or flipped to match their orders in B73Ref4. The software ALLMaps (JCVI utility libraries v1.0.6) ^63^ was conducted with default parameters, constructing 10 pseudomolecules corresponding to ten A188 chromosomes.

### BUSCO assessment

Benchmarking Universal Single-Copy Orthologs (BUSCO) ^18^ was run in a mode of “genome” to assess the completeness of the assembly with default parameters. BUSCO was run in a mode of “transcriptome” to assess the completeness of the gene annotation with default parameters. Both assessments using the Liliopsida database (liliopsida_odb10) that consisted of 3,278 conserve core genes.

### Estimation of base errors using KAD analysis

The module “KADprofile.pl” in the KAD tool (version 0.1.7) ^17^ was used to estimate errors in A188Ref1. The input read data were the merged trimmed Illumina 2×250bp reads from leaf and immature ears. The k-mer length of 47 mer was used.

### Estimation of recombination rates

Genetic distances of non-overlapping 1-Mb windows was estimated. Non-overlapping 1 Mb windows were generated by the module of “makewindows” in BEDTools (v2.29.2) ^56^. The last window of each chromosome was discarded due to the smaller size than 1 Mb. The prediction of genetic distance per window utilized a method developed previously ^64^. Briefly, a generalized additive model (GAM) was used for the prediction of the genetic distance of any physical interval.

The similar method was used to estimate recombination rates around each gene and repetitive element. For example, for a given element, we first find the midpoint of the element. The genetic positions were then predicted, by GAM, for the position 0.5 Mb upstream and the position 0.5 Mb downstream. The distance of the genetic positions was then used to represent the recombination context of the element.

The recombination rates that are smaller than 0.6 cM/Mb and higher than 3 cM/Mb were categorized to low recombination and high recombination, respectively.

### Callus induction from immature embryos

A188 ears were harvested at 11 days after pollination (DAP11), and surface-sterilized for 30 minutes in 50% (v/v) bleach (6% sodium hypochlorite) that contains 3-4 drops of Tween 20 followed by three washes in sterile distilled water. Immature embryos of size 1.0 mm-1.5 mm were isolated and cultured on callus induction medium (CIM) media ^65^. CIM was composed of Chu N6 basal medium with vitamins ^66^ supplemented with 2.3 g/L L-proline, 200 mg/L casein hydrolysate, 3% sucrose, 1 mg/L 2,4-dichlorophenoxyacetic acid, 3 g/L gelrite, pH 5.8. Subculture was conducted every 14 days. The 39-days callus samples were collected for methylome and transcriptome analysis.

### Illumina RNA-Seq, transcriptomic assembly, and differential expression

Thirty-three RNA samples were extracted from three biological replicates of each of 11 diverse tissue types of A188 with RNeasy Plant Mini Kit (Qiagen) (**Supplementary Table 10**). Briefly, the 11 tissues included the root and the above-ground of 10-day-old seedling, three different parts of the 11th leaf tissue at V12, the meiotic tassel, anther, and immature ear at V18, the endosperm and embryo 16 days after pollination, and the callus after 39 days culture of DAP11 immature embryos. RNA quality control, library preparation, and sequencing were performed on an Illumina Novaseq 6000 platform at Novogene. Trimmomatic (version 0.38) ^67^ was used to trim the adaptor sequence and low-quality bases of RNA-Seq raw reads. The parameters used for the trimming is “ILLUMINACLIP:trimming_db:3:20:10:1:true LEADING:3 TRAILING:3 SLIDINGWINDOW:4:13 MINLEN:40”. The trimming adaptor database (trimming_db) includes the sequences: adaptor1, TACACTCTTTCCCTACACGACGCTCTTCCGATCT; adaptor2, GTGACTGGAGTTCAGACGTGTGCTCTTCCGATCT. Only paired reads both of which were at least 40 bp after trimming were retained for further analysis.

Trimmed reads were aligned to A188 (A188Ref1) using HISAT2 (version 2.1.0) with the parameters of “-p 8 --dta --no-mixed --no-discordant -k 5 -x” (Kim et al., 2019). Alignments whose paired reads were concordantly paired were kept. The software StringTie2 (version 2.1.0) (Kovaka et al., 2019) was used to assemble the transcriptome with alignments from a dataset of each A188 sample with the default parameters. In total, 33 transcriptome assemblies from 33 samples were generated. All transcriptome assemblies were merged to build an A188 Illumina transcriptome assembly with the merge function in StringTie2.

### Differential expression of the callus relative to other tissue types

Trimmed reads were aligned to A188Ref1 with STAR (2.7.3a) ^68^. Uniquely mapped reads with at least 96% coverage and 96% identity were used for determining read counts per gene. DESeq2 (version 1.26.0) was used to identify differential expression between the callus and each of other tissue types. Multiple tests were corrected with the FDR (false discovery rate) approach ^69^. The FDR of 5% was set as the threshold.

### Nanopore A188 cDNA direct sequencing

Three biological replicates of the seedling and callus samples from the same tissue samples used for Illumina RNA-Seq were sequenced by the Nanopore direct cDNA sequencing protocol. Briefly, mRNA was first isolated from 10 ug total RNA with Poly(A) RNA Selection Kit (Lexogen), followed by direct cDNA library preparation with SQK-DCS109 kit (Oxford Nanopore). The protocol version for library preparation was DCS_9090_v109_revB_04Feb2019. The cDNA library was loaded onto a FLO-MIN106D R9 flowcell and sequenced on MinION (Oxford Nanopore).

FAST5 raw data was converted to FASTQ data using the basecaller Guppy version 3.4.5 (Oxford Nanopore) with default parameters. Two steps of trimming were employed. Adapter sequence was first trimmed by porechop (version 0.2.4) (https://github.com/rrwick/Porechop) with parameters “--check_reads 10000 --adapter_threshold 100 --end_size 100 --min_trim_size 5 -- end_threshold 80 --extra_end_trim 1 --middle_threshold 100 --extra_middle_trim_good_side 5 -- extra_middle_trim_bad_side 50”, and then poly A was trimmed by the software cutadapt (version 2.6) ^70^ with the options of “ -g T{12} -e 0.1 -a A{12} -n 100”. Trimmed reads were aligned to A188Ref1 as unstranded spliced long reads using MiniMap2 (version 2.14) ^71^ with the parameter “- ax splice”. Merged alignments from three replicates were input to StingTie2 for generating assembled transcripts.

### Genome annotation

The Maker (2.31.10) was used for genome annotation ^72^. The genome was masked by using Repeatmasker (4.0.7) ^73^ with the A188 repeat library built by the Extensive *de novo* TE Annotator (EDTA, v1.8.4) ^74^. Two rounds of the maker prediction were performed. At the first round, the A188 assembled transcripts and B73Ref4 protein data were used as EST and protein evidence, respectively. The parameters "est2genome=1" and "protein2genome=1" were set to directly produce gene models from transcripts and proteins. At this round, no *ab initio* gene predictors were used. Prior to the second maker round, a snap model was trained using the confident gene set from the first round. Gene models produced from round 1 were input as one of predicted gene models. These gene models were competed with gene models predicted by three gene predictors: snap (2013_11_29) ^75^, augustus (3.3.3) ^76^, and fgenesh (v.8.0.0) (softberry.com). ESTs from relative maize genotypes and proteins from closely related species were provided as additional evidence. Gene models output from Maker were further filtered. First genes matched with the following criteria “-evalue 1e-50 -qcov_hsp_perc 60” to the transposon database in Maker were filtered. Second, a transcript retained if it carried Pfam domains from the result of InterProScan (version 5.39-77.0) and/or had an annotation edit distance (AED) less than 0.4, which measured the level of discrepancy of an annotation from supporting evidence.

### Functional annotation of transcripts

BLASTP was used to map all proteins to the SWISS-Prot database (https://www.uniprot.org/) with the e-value cutoff of 1e-6. Gene ontology (GO) was extracted from InterProScan.

### Identification of a major transcript per gene

For a gene containing multiple transcripts, a major transcript per gene was selected if a transcript had the highest non-zero FPKM (Fragments Per Kilobase of transcript per Million mapped reads) determined from Illumina RNA-Seq datasets of diverse tissues by Cufflink (v2.2.1) ^77^, and/or the lowest BLASTP e-value to the SWISS-Prot database, and/or the longest transcript length. The BLASTP e-value had a priority relative to the transcript length. If data were not sufficient to make a decision, the one with the longest length was selected.

### Syntenic genes between A188 and B73

Syntenic genes were identified with MCscan (JCVI utility libraries v1.0.6) ^78^. Major transcripts were used as the input and the parameter “--cscore=.99” was used to find 1-to-1 syntenic gene relationship.

### Paralogs in A188 and orthologs between A188 and B73

Paralogs in A188 and orthologs between A188 and B73 were identified with OrthoMCL ^79^. Briefly, protein sequences of major transcripts with at least 20 amino acids were used for all-to-all BLASTP with the e-value cutoff of 1e-5. The BLASTP result was input to OrthoMCL to identify paralogous and orthologous groups.

### Identification of gene clusters

A gene cluster was defined if at least three genes from a group of A188 paralogs identified by OrthoMCL were physically closely located on a chromosome. The maximum distance is 250 kb for two neighboring genes in a cluster.

### Annotation of NLR genes

The NLR genes of A188Ref1 were annotated using the NLR-Annotator pipeline ^80^.

### Repeat annotation

EDTA (v1.8.4) ^74^ was used for repeat annotation with, maize as the “species” input, the curated maize transposable element database from https://github.com/oushujun/MTEC as the “curatedlib” input, and B73 coding sequences as the “cds’ input.

### Analysis of NUMT and NUPT

The “nucmer” command from the software MUMmer 4 ^81^ was used to align the A188 mitochondrion or chloroplast genomes to A188Ref1. For mitochondrial alignments, each required at least 5 kb and 95% identity. For chloroplast DNA alignments, each required at least 3 kb and 95% identity based on the minimal requirement for a sufficient FISH signal ^21^. Multiple alignments with the distance less than 100 kb were clustered into a block, considered to be a nuclear integration event.

### Comparative genomic analysis *via* SyRI and CGRD

The “nucmer” command was used for whole genome alignment of 10 chromosomal pseudomolecules between A188Ref1 and B73Ref4. The parameter of “--maxmatch -c 500 -b 500 -l 50” was used in the command “nucmer” and the parameter of “-i 95 -l 1000 -m” in the command of delta-filter, which resulted in best alignments with at least 1 kb matches and at least 95% identity between the two assembled genomes. The “show-coords” command with the parameter of “-THrd” was run to convert alignments to a tab-delimited flat text format. Alignment results were then used for identifying genomic structural variation and nucleotide polymorphisms through SyRI (v1.2) ^27^ with the parameter of “--allow-offset 100”. Syri analysis discovered genome duplication, translocation, inversion, as well as syntenic, unaligned, divergent sequences. SNPs, small insertions, and deletions were identified as well.

The CGRD pipeline (v0.1) (github.com/liu3zhenlab/CGRD) was employed to find copy number variation (CNV) through comparing depths of Illumina reads from A188 and B73 with the default parameters ^28^. A value of the log2 read depth ratio per sequence segment (LogRD) is the indication for CNV. For a segment, the LogRD is close to zero if sequences of two genotypes are identical and no CNVs. The sufficient derivation of the mean of LogRD from zero is likely due to CNV or a high level of divergence. CGRD was performed using A188Ref1 as the reference genome and identified sequences of A188Ref1 showing conserved (B73=A188), copy number plus (B73>A188), and copy number minus (B73<A188) in B73 relative to A188. When B73Ref4 was used as the reference, the analysis found sequences of B73Ref4 showing conserved (A188=B73), copy number plus (A188>B73), and copy number minus (A188<B73) in A188 relative to B73.

### Identification of PAV and highly divergent sequences (HDS)

SyRI analysis listed B73Ref4 sequences that were not aligned to A188Ref1, and *vice versa*, as well as insertion/deletion polymorphisms between the two chromosomal sequences. Unaligned sequences or insertion/deletion polymorphisms identified by SyRI were compared with CGRD segments. For each SyRI event, a supporting score of read depth data from CGRD was determined by using the formula of 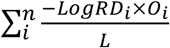, where *i*; represents the *i*th overlap between a CGRD segment and a SyRI event; *LogRD*; stands for the *LogRD* of the CGRD segment and only negative values were taken into calculation; *O* is the overlapping length in bp; *L* is the length in bp of the SyRI event; and *n* is the total number of overlaps. The resulting value from the formula represents the degree of the differentiation in read depth between the two genotypes for the SyRI event. The higher the number, the more confidence the SyRI event is a PAV or HDS. A SyRI event is considered to be a PAV or HDS if a supporting score is larger than 3.

### Identification of large inversion events

Inversion, from kb to Mb levels of events, between A188Ref1 and B73Ref4 were revealed by SyRI. Large events with both A188 and B73 sequences larger than 0.5 Mb were exacted. First, the inversion sequences of B73Ref4 were aligned to B73Ref5 to confirm the inverted orientation relative to A188Ref1. For a given inversion, if >80% B73Ref4 sequences were aligned to B73Ref5 in the plus orientation, the inversion was supported by B73Ref5. If <20% B73Ref4 sequences aligned to B73Ref5 were in the plus orientation, the inversion was considered to not be supported by B73Ref5. Second, the recombination frequency between the start and the end of an inversion event was estimated and adjusted to cM per Mb. Third, SNPs between the two genomes and located on the inversion were identified. The common SNPs genotyped in the maize 282 ^82^ population were extracted for determining linkage disequilibrium (LD) between SNPs in distance of 0.2-0.3 Mb. Vcftools (v0.1.17) ^83^ was employed to calculate LD. The genome-wide LDs between SNPs in distance of 0.2 Mb were determined as the control.

### Structure analysis of inversions in maize HapMap2 population

The software STRUCTURE (v2.3.4) ^84^ was used to analyze the inference of population structure for A188 inversions in maize HapMap2 population ^85^. A188 and B73 SNPs between inversion regions were discovered by SyRI. HapMap2 genotyping data overlapping with inversion SNPs were extracted and the subset SNPs with the missing rate less than 20% were input for STRUCTURE analysis. The major alleles, minor allele and missing locus in SNP dataset were converted to 0, 1, and −1, respectively. K=2 as the cluster number and 10 replicate runs of the admixture model were used, with a burn-in of 10,000 iterations and a run length of 20,000 steps.

### Fluorescence *in situ* hybridization (FISH)

Mitotic and meiotic chromosomes were prepared as described by Koo and Jiang (2009) with minor modifications ^86^. Root tips were collected from seedling plants and treated in a nitrous oxide gas chamber for 1.5 h, fixed overnight in ethanol:glacial acetic acid (3:1), and then squashed in a drop of 45% acetic acid. Anthers were squashed in 45% acetic acid on a slide and checked under a phase microscope. All preparations were stored at −70°C until use.

DNA probes of the CentC, Knob, Cent4 ^87^, and the probes for examining NUMTs, the PME cluster, and a potential large inversion on chromosome 4 (**Supplementary Table 11**) were labeled with digoxigenin-11-dUTP (Roche, Indianapolis, IN), biotin-16-dUTP (Roche), and/or DNP-11-dUTP (PerkinElmer), depending on whether two or three probes were used in the FISH experiment ^87^. The FISH hybridization procedure was according to a previously published protocol ^88^. After post-hybridization washes, the probes were detected with Alexafluor 488 streptavidin (Invitrogen) for biotin-labeled probes, and rhodamine-conjugated anti-digoxigenin for dig-labeled probe (Roche). The DNP-labeled probe was detected with rabbit anti-DNP, followed by amplification with a chicken anti-rabbit Alexafluor 647 antibody (Invitrogen). Chromosomes were counterstained with 4′,6-diamidino-2-phenylindole (DAPI) in Vectashield antifade solution (Vector Laboratories). The images were captured with a Zeiss Axioplan 2 microscope (Carl Zeiss Microscopy LLC) using a cooled CCD camera CoolSNAP HQ2 (Photometrics) and AxioVision 4.8 software. The final contrast of the images was processed using Adobe Photoshop CS5 software (Adobe).

### QTL mapping

Kernel colors of 125 DH lines were scored as 1 to 6 (1=white, 6=yellow, and 2 to 5 indicated colors between white and yellow). QTL mapping of kernel color was performed by using scanone function with Haley-Knott regression method in the R package R/qtl ^89^. The LOD cutoff was the 5% highest LOD value from 1,000 permutations of phenotypic data.

### qRT-PCR

qRT-PCR was used to measure the gene expression of *ccd1* and *y1* gene in genotypes of A188, B73, DH305 and DH312. Immature ears of the four genotypes were harvested from the summer nursery in Manhattan Kansas at 16 days after pollination (DAP16). Fifteen kernels were randomly sampled from the middle of an ear, five kernels of which were pooled as a biological replication for RNA isolation. cDNA was synthesized with Verso cDNA Kit (Thermo Scientific) following the manufacturer’s protocol. qRT-PCR was performed in a reaction of 10 ul with the IQTM SYBR Green Supermix reagent (BioRad) on the CFX96 Real-Time PCR System (BioRad). The thermocycling conditions for the PCR are initial denaturation at 95°C for 3 minutes, followed by 40 cycles of denature at 95°C for 15 seconds, annealing and extension at 60°C for 30 seconds. The housekeeping reference gene *actin1* was used as the internal control. Cycle threshold values (*Ct*) of two technical replicates were averaged and used to quantify relative gene expression. The relative expression of each of *ccd1* and *y1* genes in each sample was calculated using the formula 100 × 2^*actinCT−geneCt*^, where *actinCt* and *geneCt* stand for the *Ct* values of *actin1* and *ccd1* (or *y1*), respectively. The primers used are as follows: *actin1*: act1_qrt_2F and act1_qrt_2R; *ccd1*: ccd1_qrt_5F and ccd1_qrt_5R; *y1*: y1_qrt_4F and y1_qrt_4R. Sequences of primers are in **Supplementary Table 11**.

### Whole-genome bisulfite sequencing (WGBS)

Two biological replicates of each of two out of eleven tissue types used in RNA-Seq, the seedling and the callus, were subjected to WGBS on a Novaseq 6000 at Novogene. A Bismark pipeline (v0.22.1) was adapted to process bisulfite sequencing DNA methylation data ^90^. Briefly, raw reads were subjected to Trimmomatic trimming (v0.38) ^67^ to remove adaptor and pool-quality sequences. Bowtie2 (v2.3.5.1) ^91^ was used for the alignment and alignments of duplicated reads were removed before methylation calling. The methylation level per cytosine site of all three sequence contexts (CG, CHG, and CHH) were determined, which were used for identifying differentially methylated regions (DMRs) with the DSS R package (v2.34.0) (github.com/haowulab/DSS).

### DNA methylation around genes and on repetitive sequences

Genomic regions (gene body) from the translation start site (TSS) to the translation termination site (TTS), which were based on genomic locations of major transcripts, were equally divided into 200 windows. For each gene, the 2 kb 5’ upstream region and the 2 kb 3’ downstream region of each gene were also extracted. The DNA methylation rate in three sequence contexts (CG, CHG, and CHH) on each window of the gene body or each 20 bp in upstream and downstream regions was separately determined for examining the distribution of DNA methylation on and around genes.

DMRs were located in the three regions, 5’ upstream 1 kb, gene body, 3’ downstream 1 kb. For each region, the independence between changes of DNA methylation, increased or decreased in the callus versus the seedling, and regulation in gene expression, up- or down-regulated in the callus versus the seedling from DE analysis, was examined through χ^2^statistical test. Tests were performed for all three methylation types: CG, CHG, and CHH.

DNA methylation rate per 100 bp of repetitive sequences was determined. Annotation of repetitive types was from EDTA and additional 45S rDNA alignment analysis. Paired t-test was performed between the two tissues: callus and seedling.

### Tissue network and principal component analyses of A188 tissues

The A188 tissue network was constructed with the R package WGCNA (version 1.66) ^92^ using expression of 29,222 genes in 33 RNA-Seq datasets from 11 A188 tissue types. WGCNA was performed to clustered A188 tissue samples with the parameters minModuleSize = 6 and soft-thresholding power = 9. The Gephi software (version 0.9.2) ^93^ was used to visualize tissue networks with the module and connectivity information from the WGCNA result. Principal component analysis (PCA) was also performed using the R functions *prcomp* with the expression per gene averaged from three replicates per tissue type.

### Gene ontology (GO) enrichment analysis

The enrichment analyses were performed to determine if a certain GO was over-represented in a selected group of genes. The resampling method in GOSeq ^94^ was employed.

## Supporting information

ExtendedData_1_SyRI.out.simple.txt

ExtendedData_2_CGRD_A2B_B73Ref4.txt

ExtendedData_3_CGRD_B2A_A188Ref1.txt

ExtendedData_4_A188.PAV.HDS.genes.bed.txt

ExtendedData_5_B73.PAV.HDS.genes.bed.txt

ExtendedData_6_normalized.read.counts.txt

ExtendedData_7_CG.DMRs.all.txt

ExtendedData_8_CHG.DMRs.all.txt

ExtendedData_9_CHH.DMRs.all.txt

ExtendedData_10_BAgm.v02.txt

## ACKNOWLEDGEMENTS

We thank Drs. Candice A.C. Gardner and Mark J. Millard at the North Central Regional Plant Introduction Station for distributing A188 seeds and Dr. Candice Hirsch from University of Minnesota to provide guidance of tissue collection for RNA-Seq. We thank the Doubled Haploid Facility at Iowa State University for producing DH lines and computational support from the Beocat High-Performance Computing Facility at Kansas State University. We thank funding support from the US National Science Foundation (award No. 1741090), the USDA’s National Institute of Food and Agriculture (award no. 2018-67013-28511), and the Agricultural Science and Technology Innovation Program of Chinese Academy of Agricultural Sciences. This is the contribution number 21-040-J from the Kansas Agricultural Experiment Station.

## AUTHOR CONTRIBUTIONS

S.L., F.F.W, S.P. M.C. and H.W. conceived and designed experiments. G.L., D.K., H.Z., T.M.T., Y.L. (Yan L.), M.Z., Y.H., Y.Q., Y.L. (Yunjun L.) performed experiments and collected data. G.L., C.H., J.L., H.T., S.L. analyzed data. H.L. participated in the overall design of the strategies of genome assembly and genome comparison. B.W. assisted in the genome annotation. F.M., H.K. and S.M.K. participated in maintenance of DH lines. J.Z. and G.W. provided Bionano data. S.L. and P.S.S conceived the CGRD approach. S.L. and N.S. designed and interpreted DNA methylation experiments. D.R.M. contributes ideas and discussion on the kernel color experiment. G.L., C.H., J.Z, H.W., F.F.W., N.S., P.S.S, G.W., S.L. wrote the manuscript with comments from other authors.

## DATA AVAILABILITY

The A188Ref1 genome sequence is available at NCBI under the accession (JABWIA000000000). The annotation (A188Ref1a1) is available at MaizeGDB.org. Raw Nanopore whole genome sequencing data, Illumina whole genome sequencing data, Nanopore cDNA sequencing data, Illumina RNA-seq data, and whole-genome bisulfite are available at NCBI SRA under the project of PRJNA635654. Essential scripts related to the manuscript are available at github.com/liu3zhenlab/A188Ref1. The CGRD pipeline can be downloaded from github.com/liu3zhenlab/CGRD.

