## Supplementary material for "Chromosome-level Genome Assembly of a Regenerable Maize Inbred Line A188": A188_suppl_figures.pdf

Lin el al.

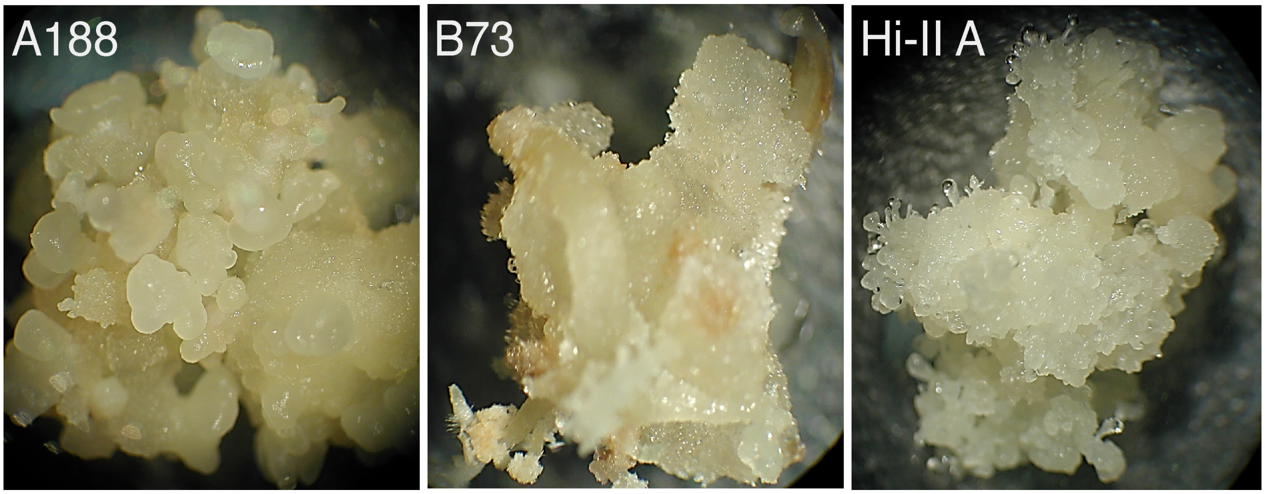


**Supplementary Figure 1. Calli from immature embryos of A188, B73, and Hi-II A. a**, **b**, **c**) White, compact, and nodulated somatic embryos and embryogenic callus from A188 **(a)**, brownish, loose and non-embryogenic callus from B73 **(b)**, and white, friable and embryogenic callus from Hi-II A (**c**) after 28 days of culture in callus induction media.

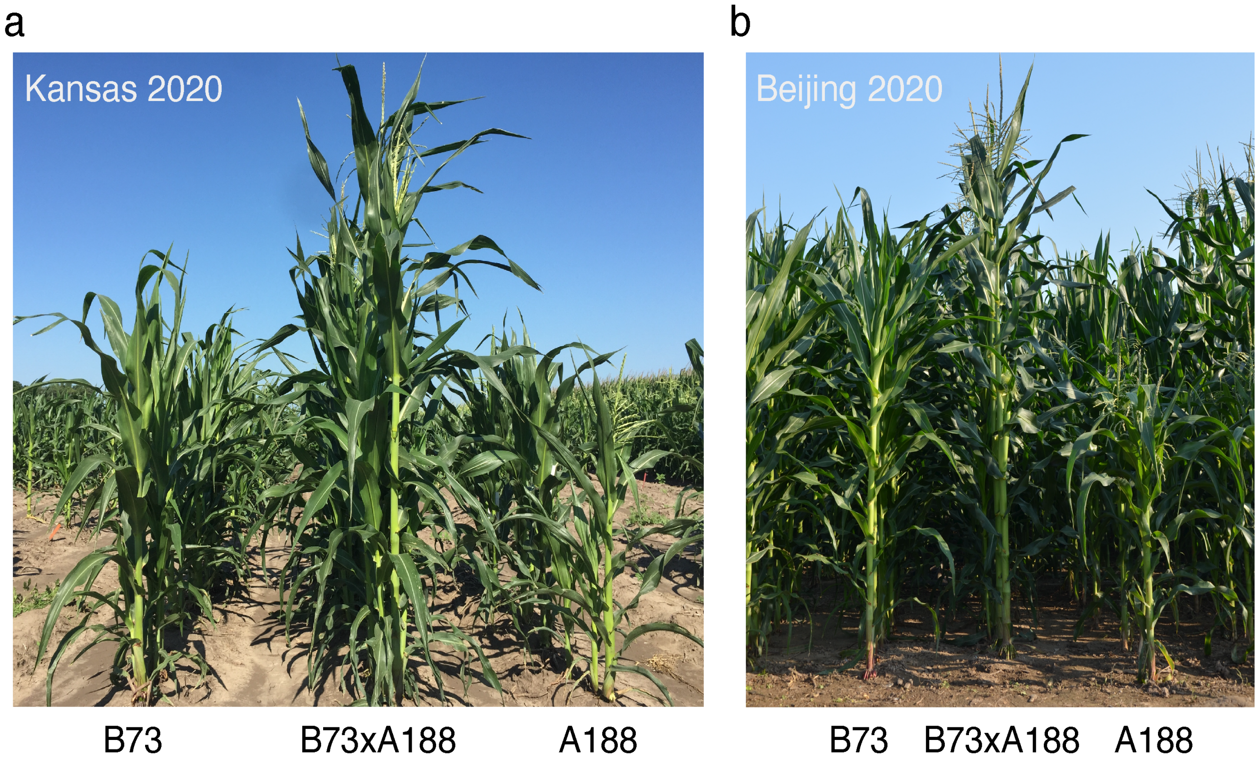


**Supplementary Figure 2. Phenotypes of B73, F1, and A188. a,b**). Plants from 2020 summer nursery in Kansas (**a**) and Beijing (**b**).

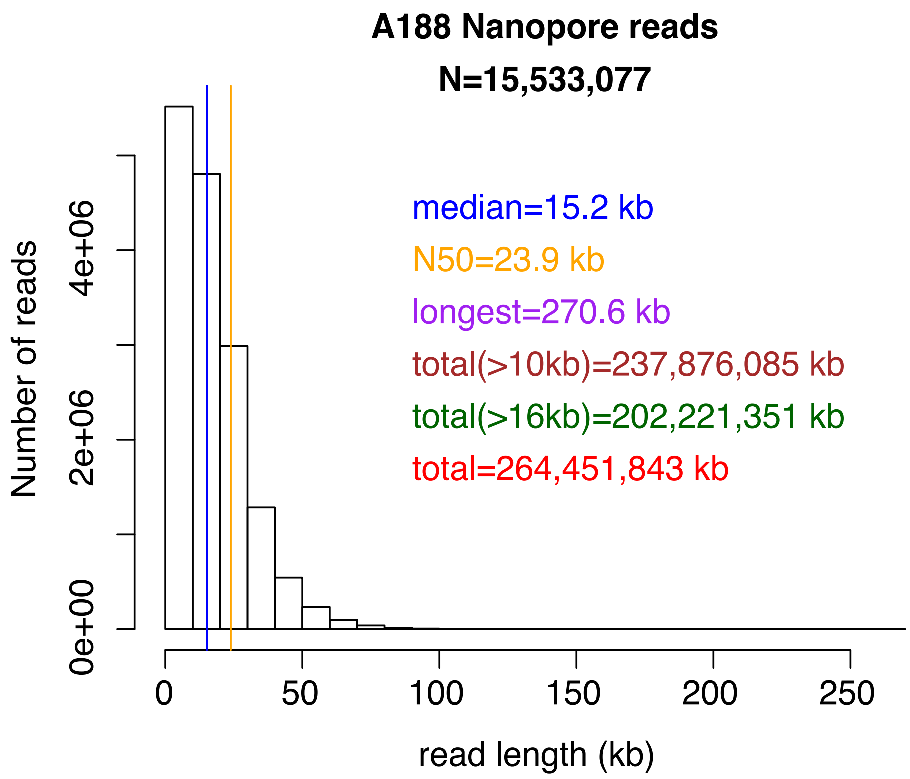


**Supplementary Figure 3. Histogram of lengths of Nanopore raw reads.** Thirty one MinION flowcells were used to produce Nanopore reads for the A188 genome assembly, producing >264 Gb total sequences. The median and N50 of read lengths are indicated by blue and red vertical lines, respectively.

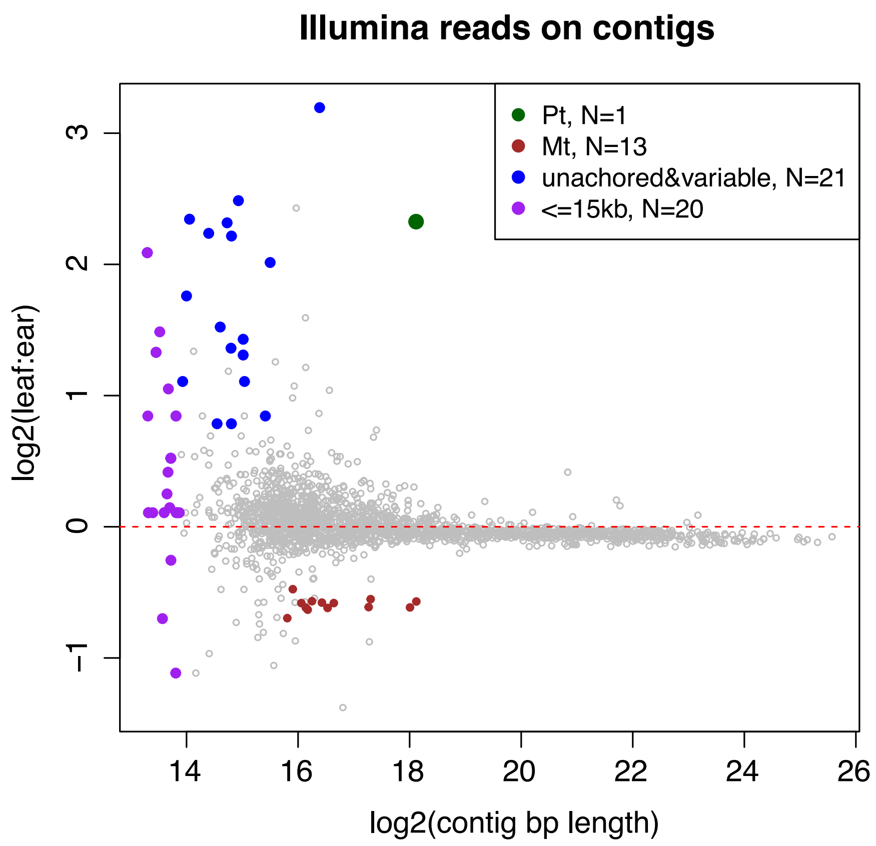


**Supplementary Figure 4. Contig filtering based on read depths.** Log2 values of Illumina read depth ratios of seedling leaf to ear samples, log2(leaf:ear), were determined for each contig. Contigs that were not anchored to B73Ref4 and showed a high variability from 0 of log2(leaf:ear) and contigs less than 15 kb were discarded. The chloroplast contig (pt) and mitochondrial contigs (mt) were replaced by the A188 chloroplast complete sequence (Genbank accession KF241980.1) and the A188 mitochondrion complete sequence (Genbank accession DQ490952.1), respectively.

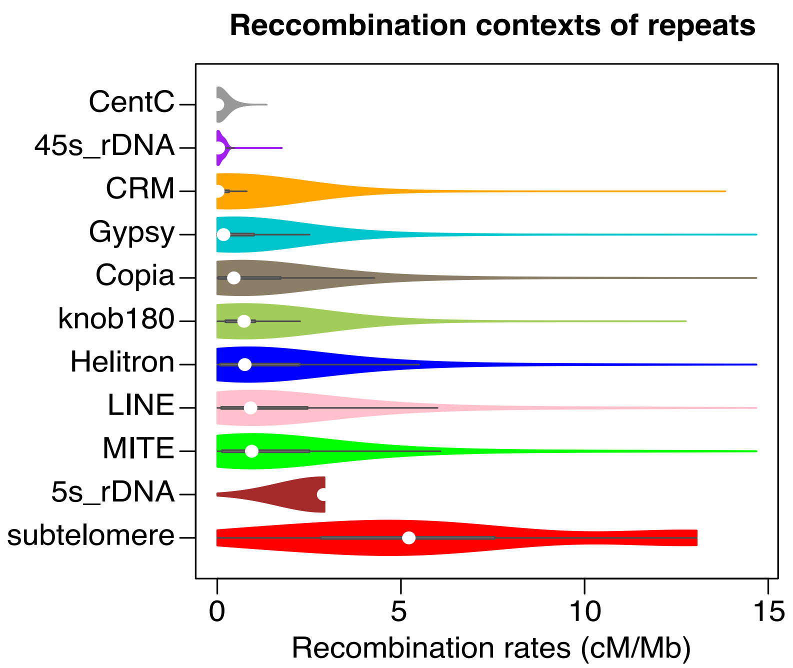


**Supplementary Figure 5. Recombination of contexts around repeats.** For each repeat type, the recombination rate (cM/Mb) of the surrounding 1 Mb context was estimated for each sequence on chromosomes. A violin plot of all sequences of each type was plotted with a dot to represent the median.

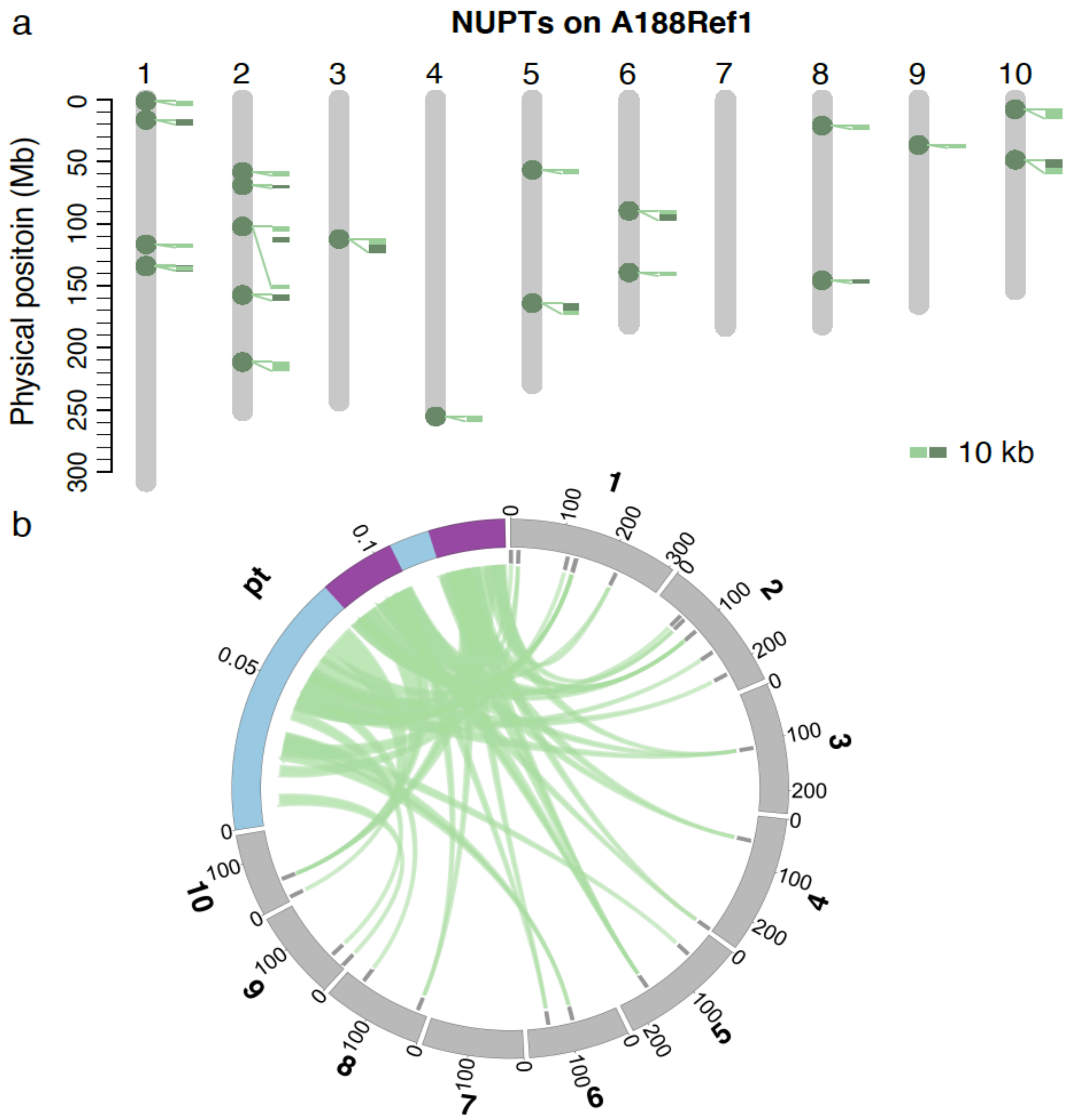


**Supplementary Figure 6. Nuclear integration of chloroplast DNA. a**). NUPT sequence on 10 chromosomes of A188Ref1. Each dot on chromosomes designates a potential NUPT integration. Close-up alignments with the chloroplast genome are shown along NUPTs. Each alignment requires at least 3 Kb match and 95% identity. **b**). Circos plot of alignments between the chloroplast (pt) genome and ten chromosomes. Purple highlighted the large duplicated regions on pt. Gray bars locate NUPT positions. Note that the chromosomal scale is different from the pt scale. Numbers on the track are in Mb.

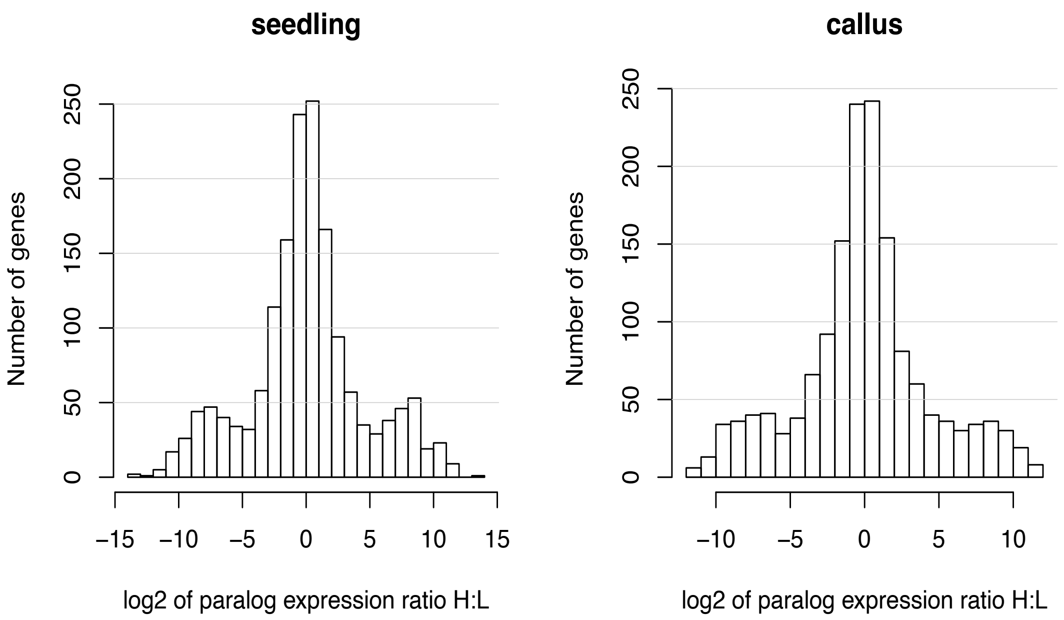


**Supplementary Figure 7. Expression comparison of paralogs in high- and low-recombination regions.** Pairs of paralogs of which one is located at a high-recombination region (H) and the other is located at a low-recombination region (L) were compared for their expression. Histograms of the log2 values of read counts ratio of H to L were plotted. Relatively symmetric distributions between positive and negative log2 values indicated that the genomic context of the gene location was not a major driver for gene expression.

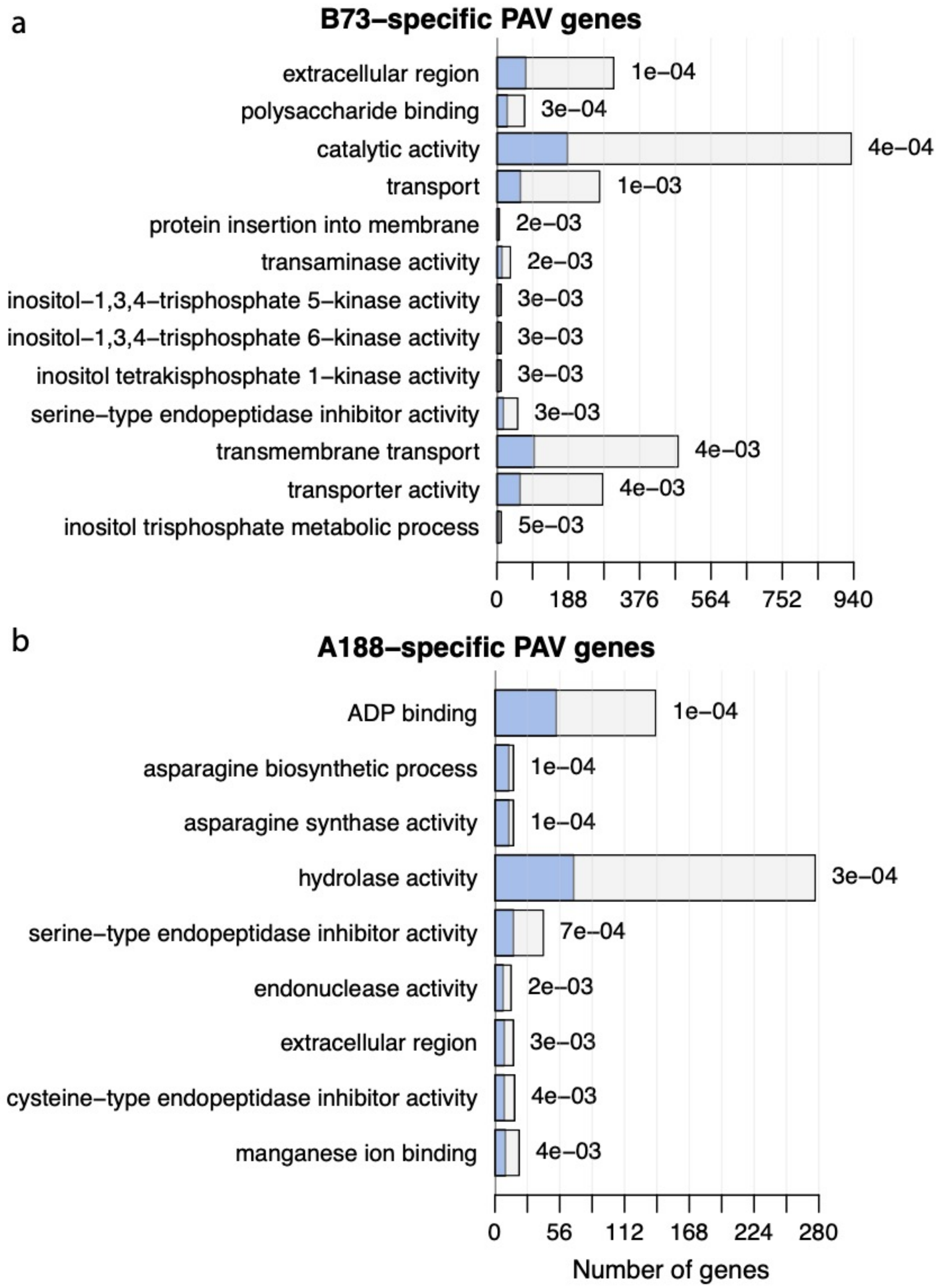


**Supplementary Figure 8. GO enrichments of PAV/HDS genes.** Enriched GO terms in B73-specific PAV/HDS genes (**a**) and in A188-specific PAV/HDS genes (**b**). In each barplot, a blue bar stands for the number genes in the PAV/HDS gene set and the whole bar (blue and empty) stands for the total number of genes of the associated GO term. P-values were labeled on the top of each bar. Only the GO terms with the p-value smaller than 0.005 and containing at least five PAV genes were plotted.

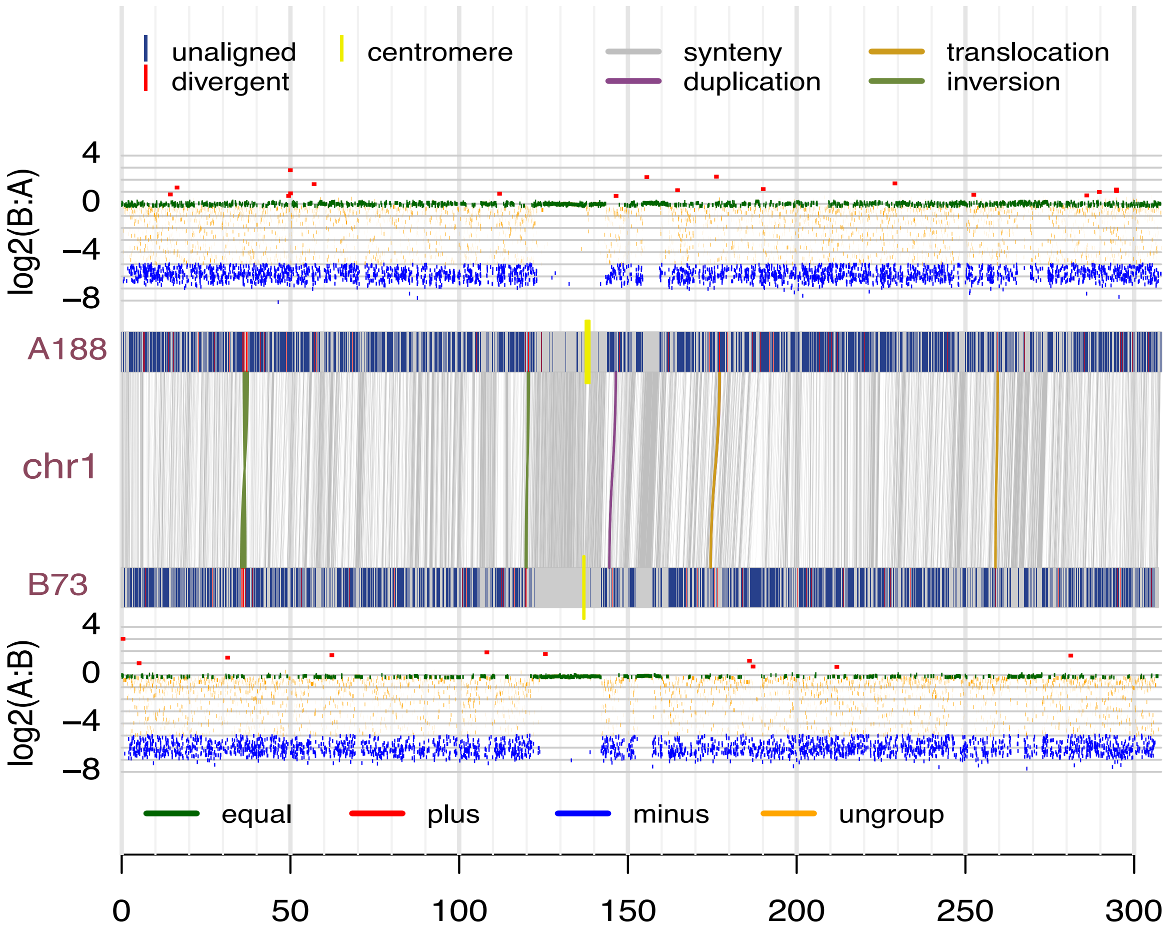


**Supplementary Figure 9. SyRI and CGRD results on chromosome 1.** CGRD results using A188Ref1 and B73Ref4 as the reference genomes were plotted on the top and bottom, respectively. Y-axis represents log2 values of ratios of read depths of B73 to A188, log2(B:A), or log2 values of ratios of read depths of A188 to B73, log2(A:B), signifying copy number variation (CNV). The SyRI result is displayed in between two CGRD results. Alignments of syntenic blocks larger than 10 Kb and alignments of other rearrangements larger than 0.5 Mb are plotted. On each A188 and B73 chromosome, segments not aligned to the other genome (unaligned), segments divergent with the other genome in a high degree (divergent), and centromeres are highlighted. The same plotting strategy was applied to chromosome 2, 3, 5, 6, 7, 8, 9, and 10.

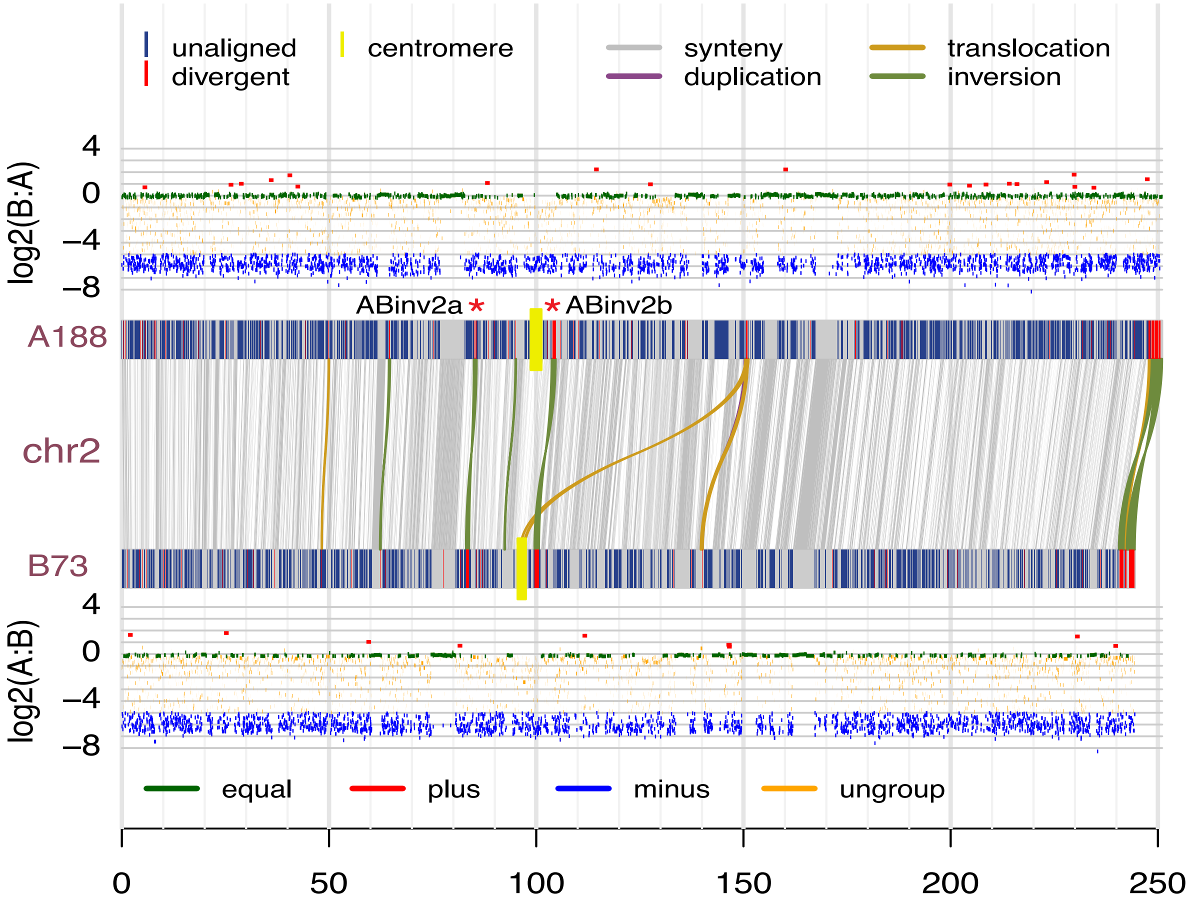


**Supplementary Figure 10. SyRI and CGRD results on chromosome 2.** The red * labels a well-evidenced inversion.

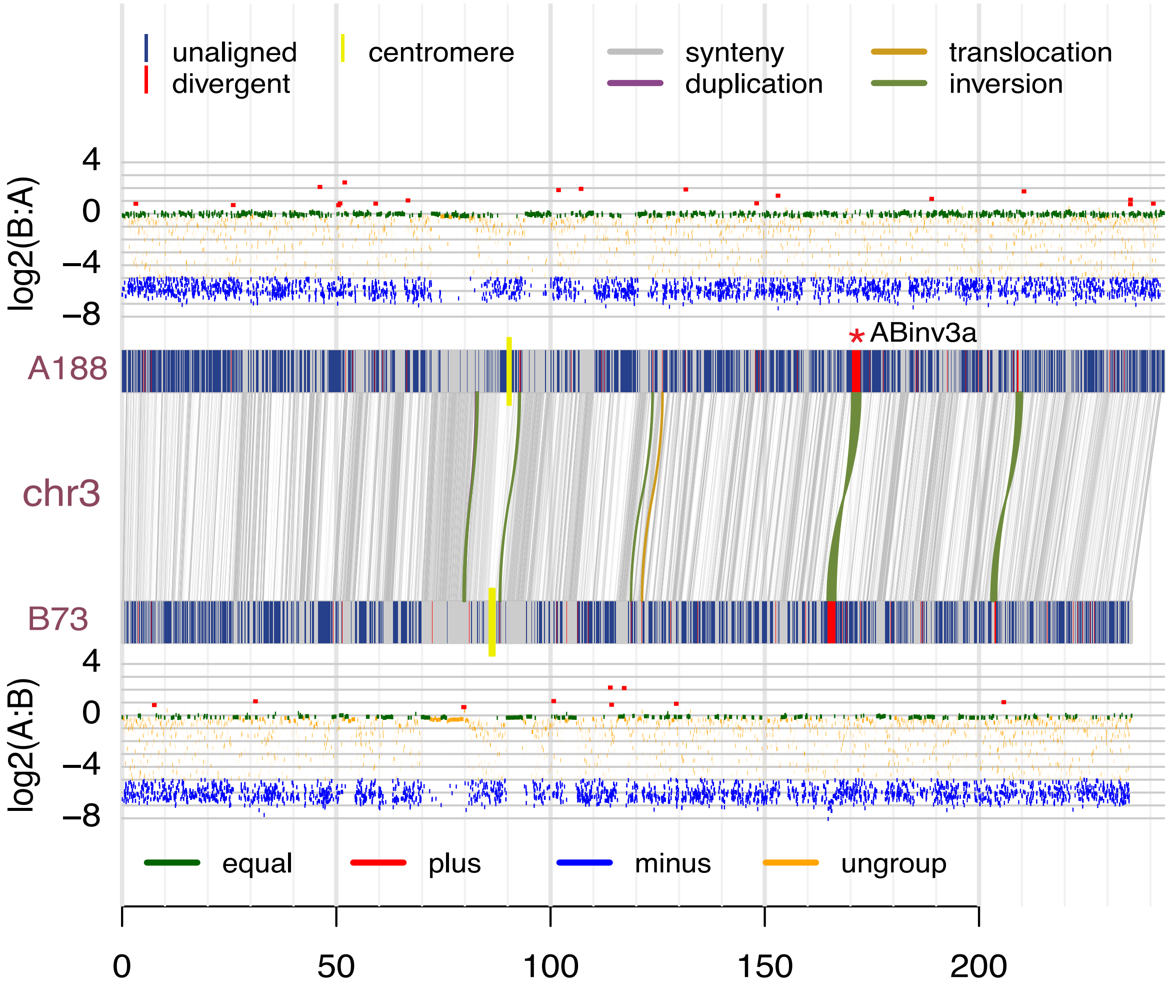


**Supplementary Figure 11. SyRI and CGRD results on chromosome 3.** The red * labels a well-evidenced inversion.

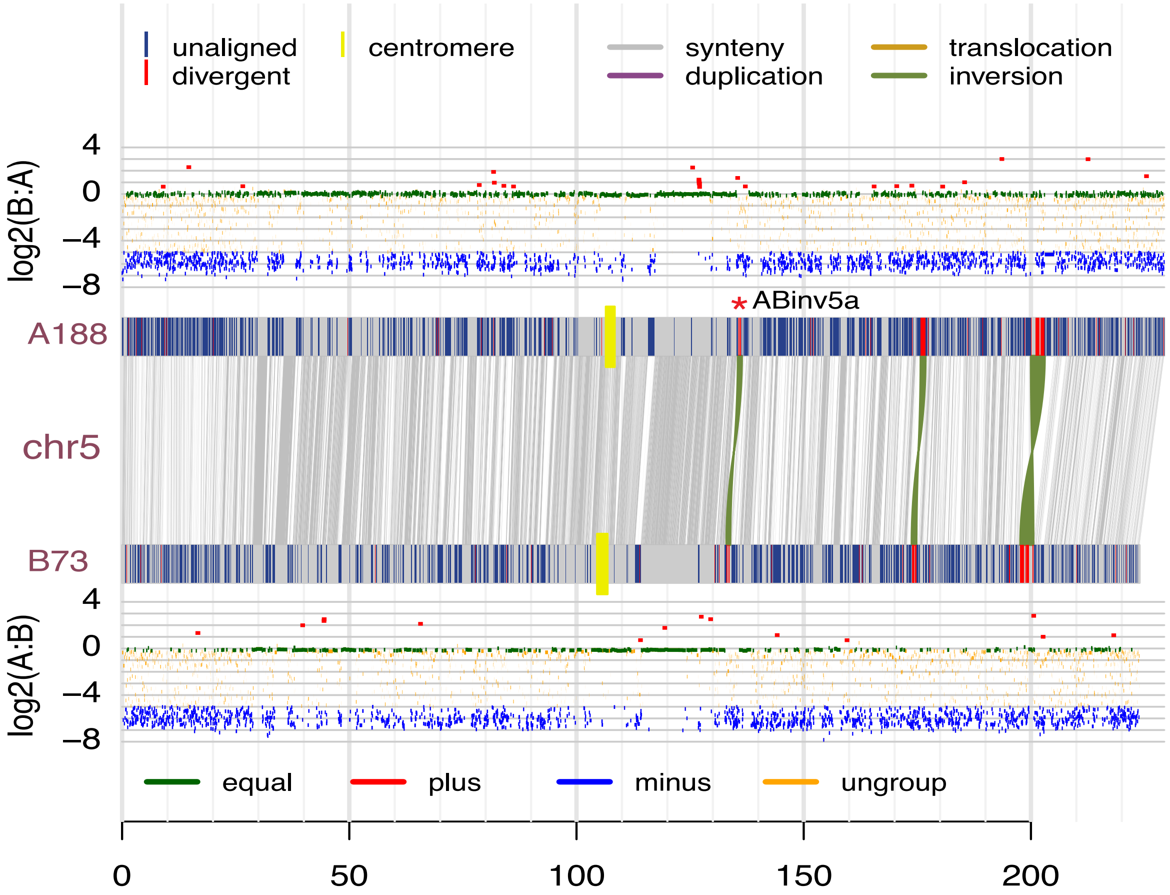


**Supplementary Figure 12. SyRI and CGRD results on chromosome 5.** The red * labels a well-evidenced inversion.

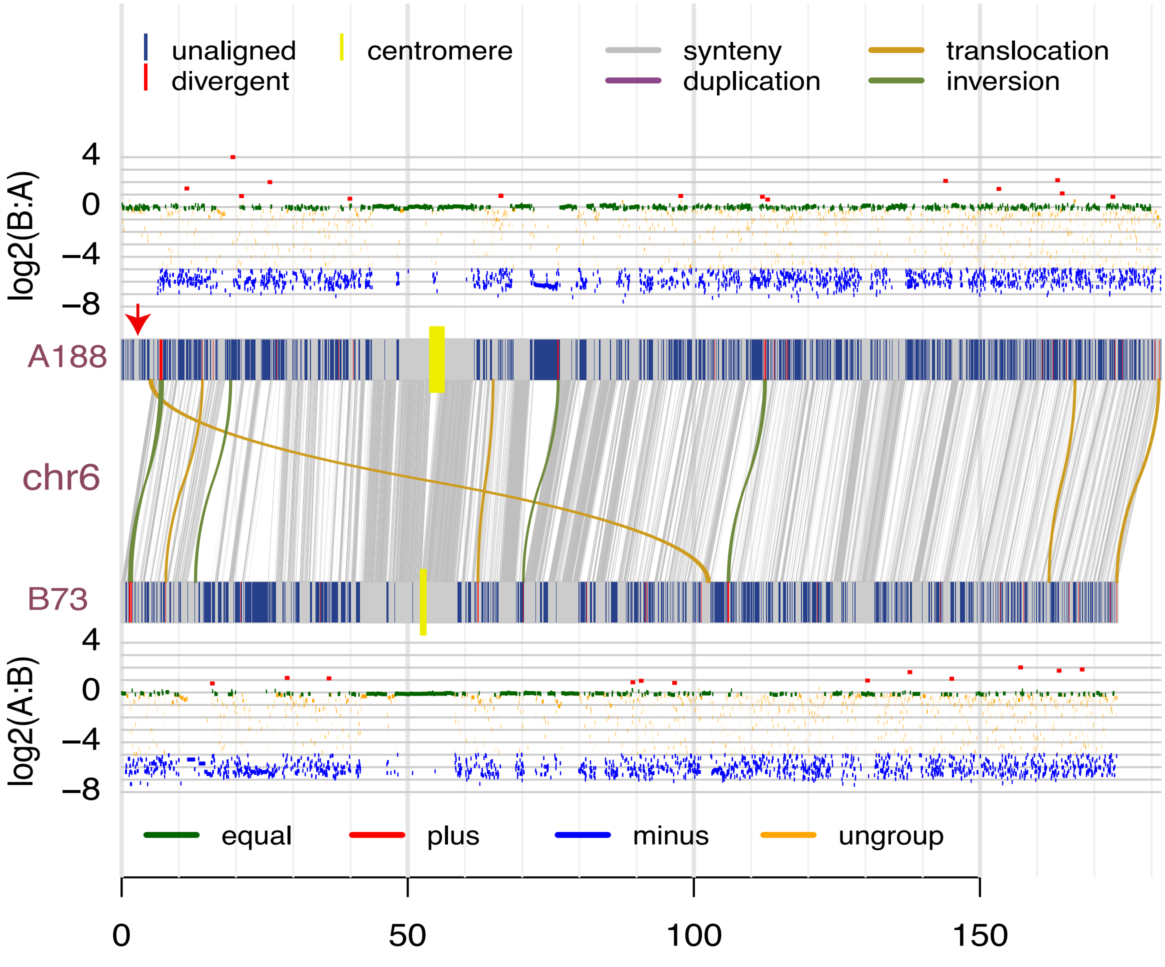


**Supplementary Figure 13. SyRI and CGRD results on chromosome 6.** The arrow points at a relative conserved region (equal) between the two genomes. However, the region is missed in the B73Ref4. The newly assembled B73Ref5 has the region at the beginning of chromosome 6.

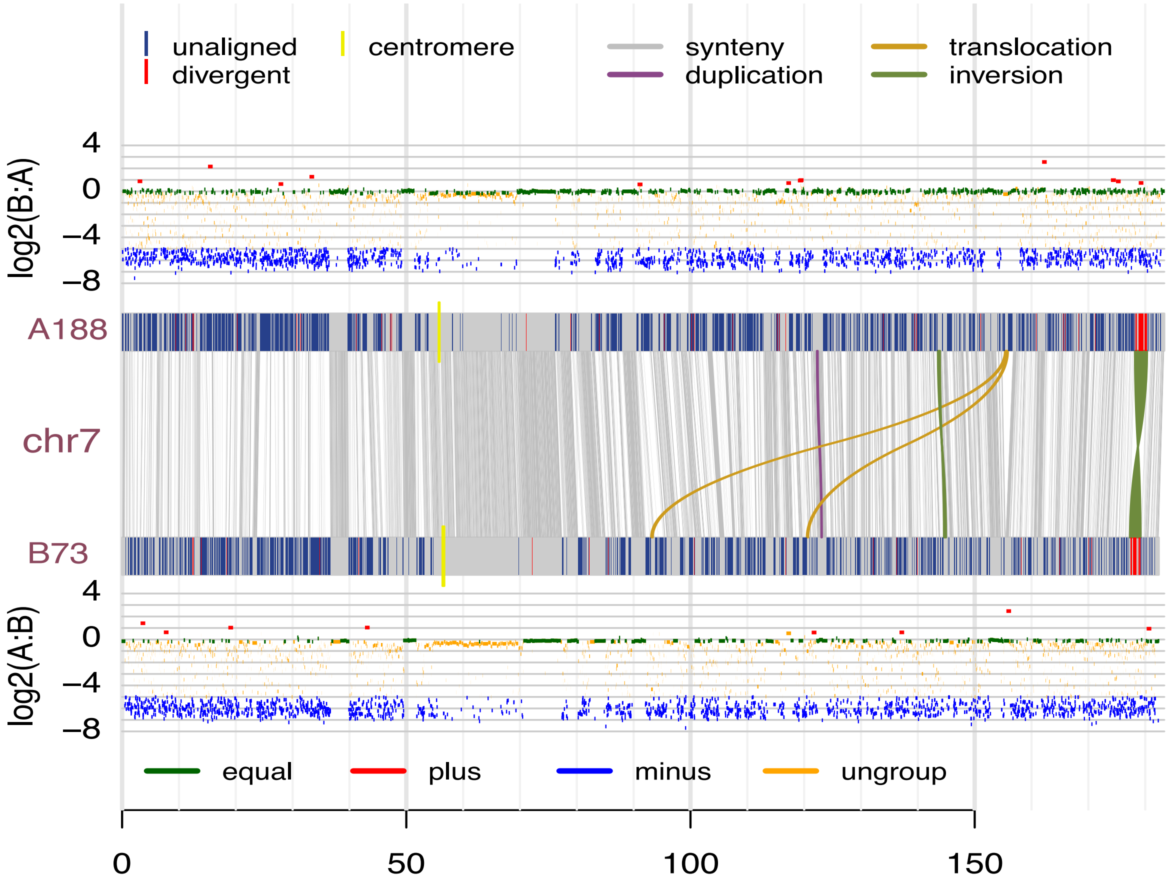


**Supplementary Figure 14. SyRI and CGRD results on chromosome 7.**

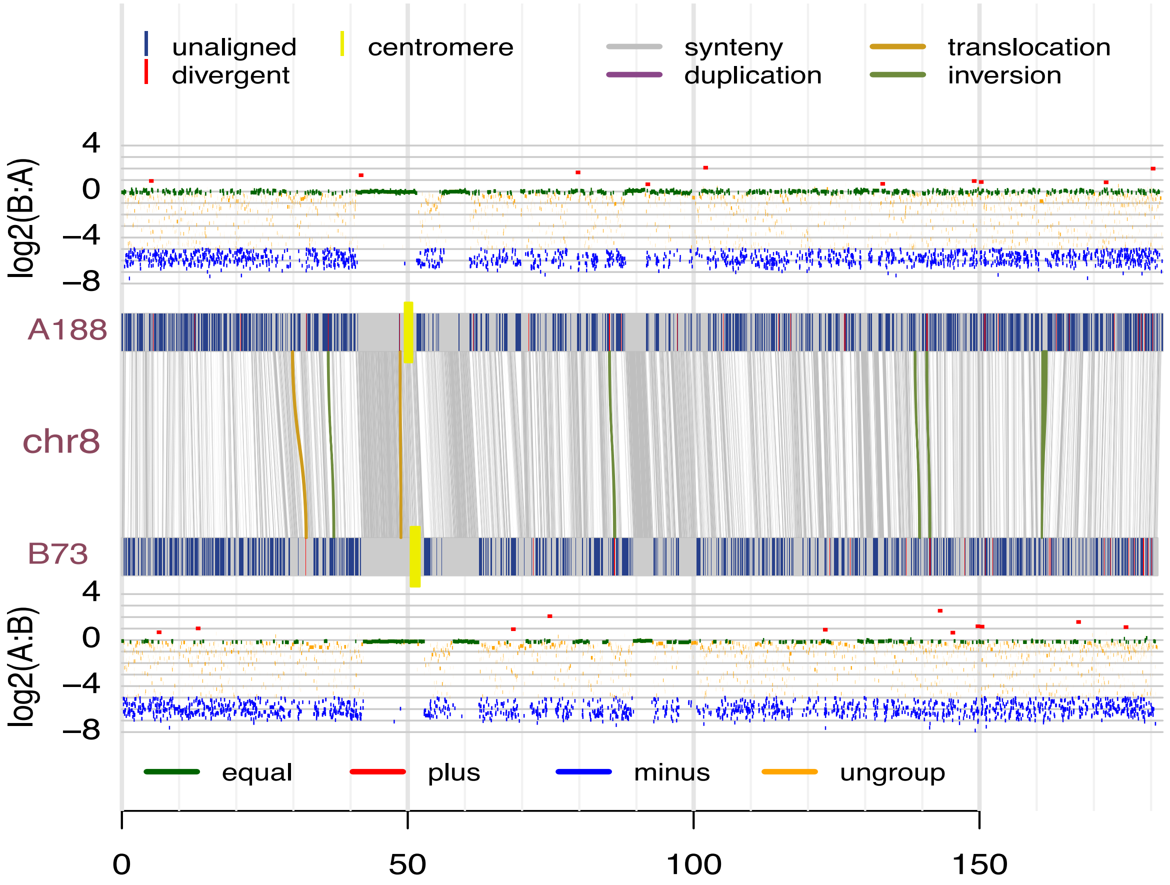


**Supplementary Figure 15. SyRI and CGRD results on chromosome 8.**

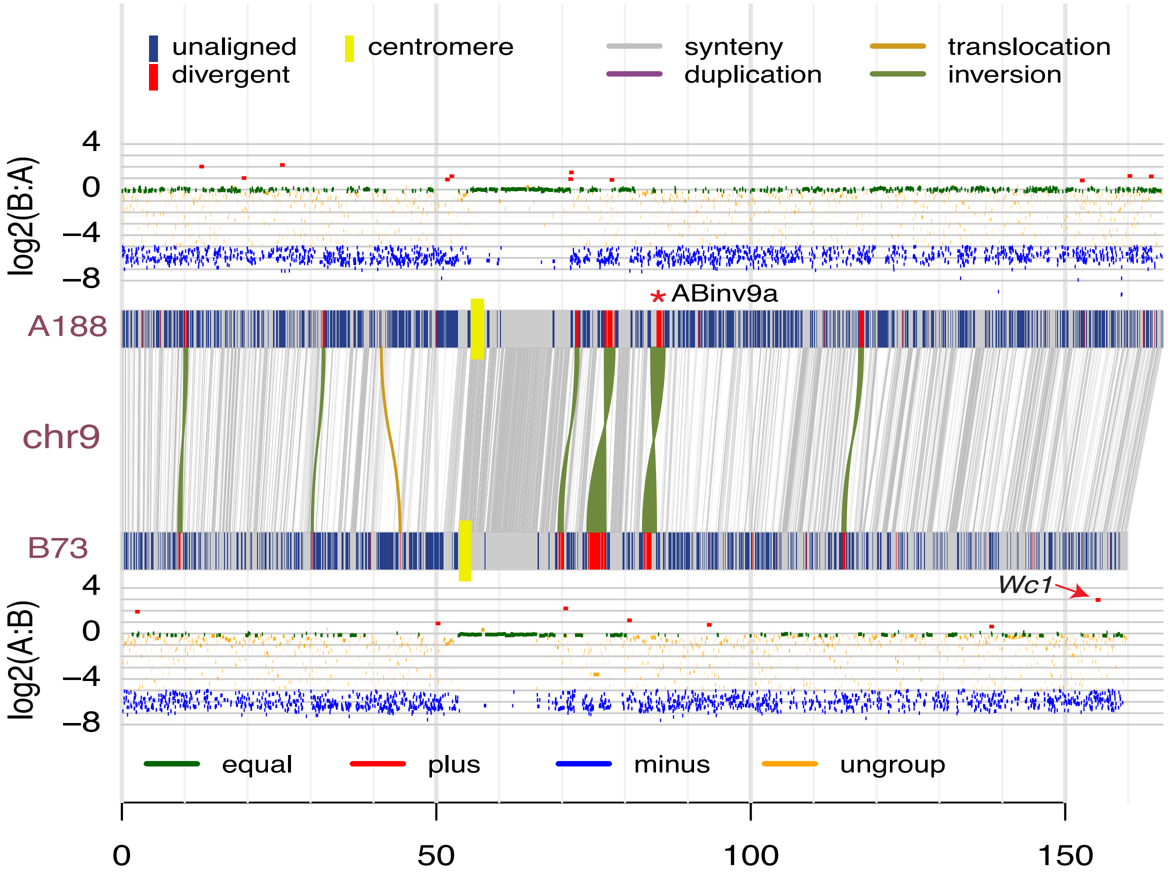


**Supplementary Figure 16. SyRI and CGRD results on chromosome 9.** The red * labels a well-evidenced inversion. The arrow points at the B73 *Wc1* region showing A188plus, which A188 has higher copy number as compared to B73.

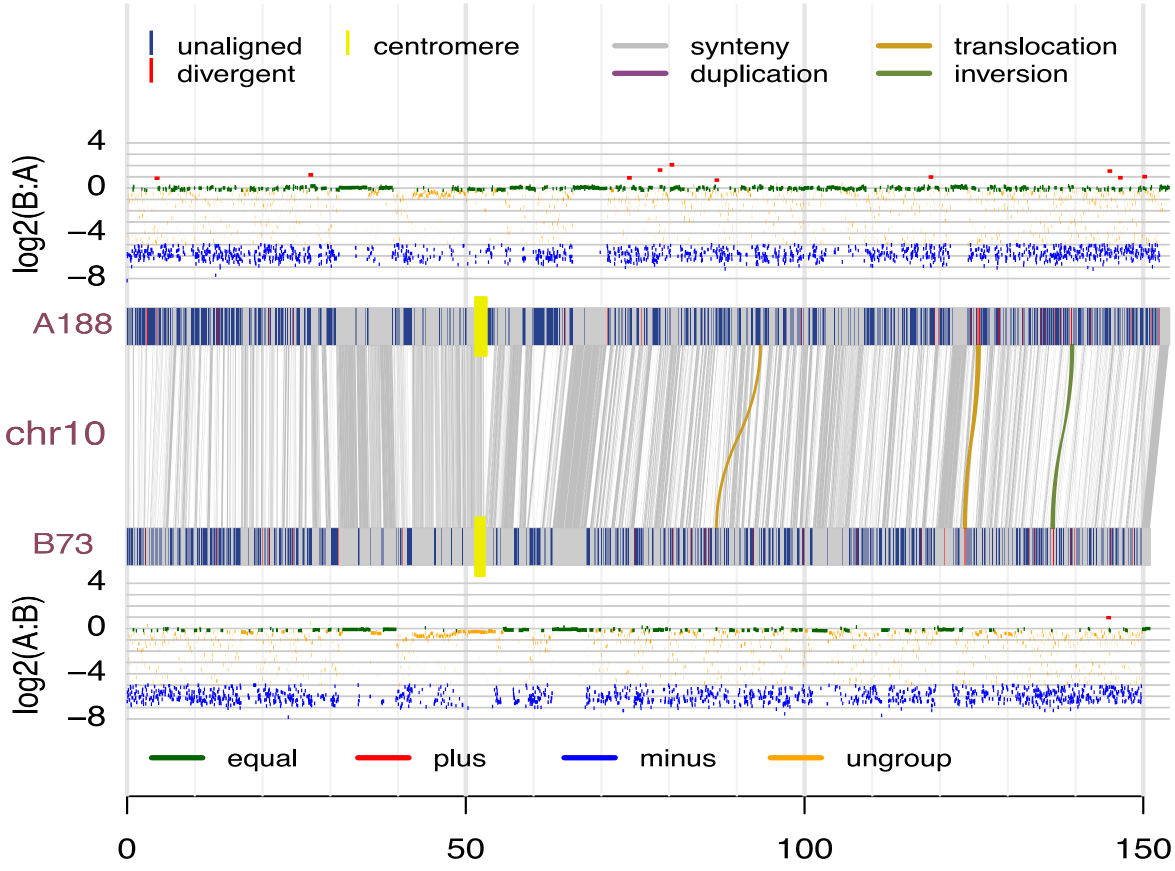


**Supplementary Figure 17. SyRI and CGRD results on chromosome 10.**

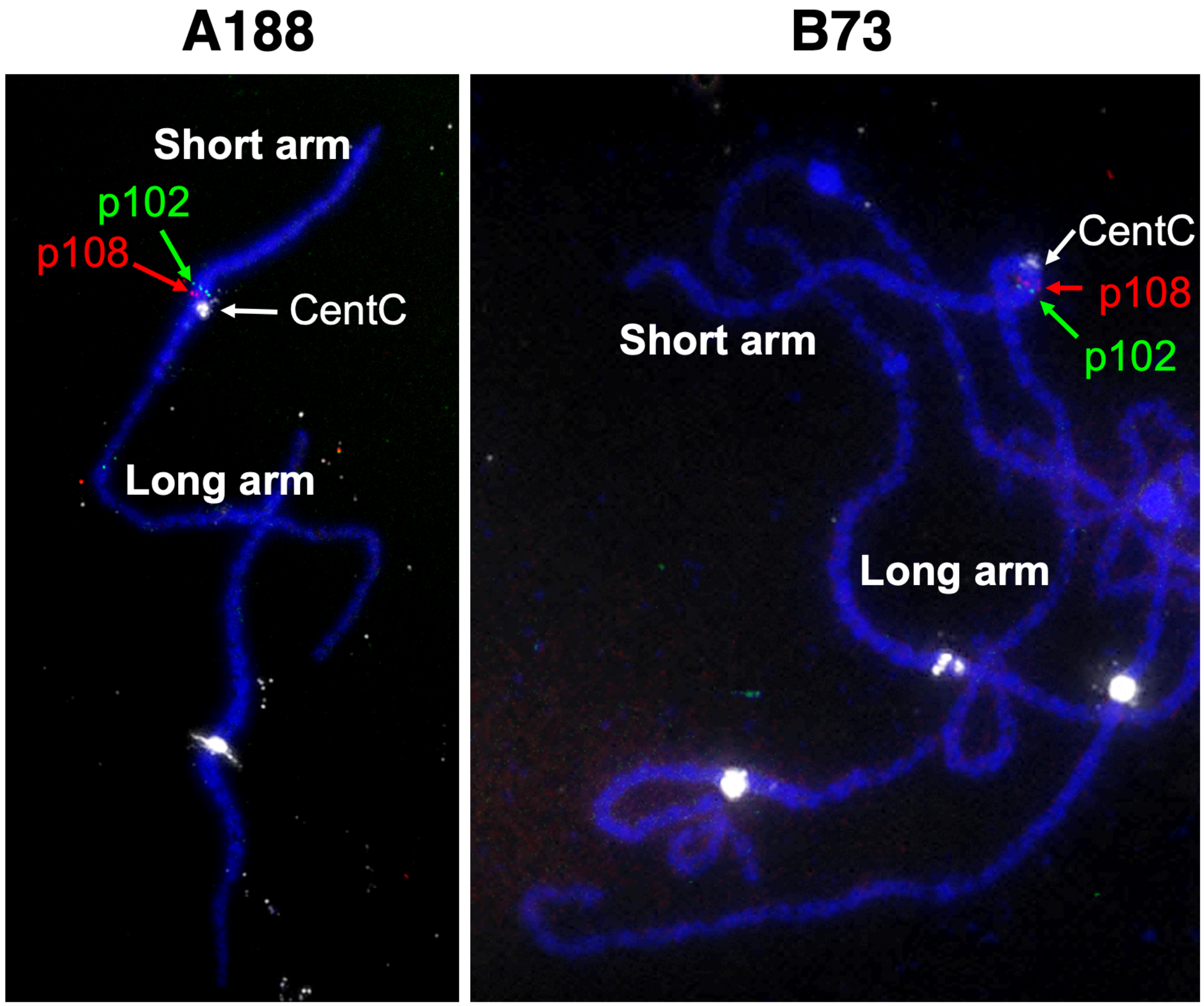


**Supplementary Figure 18. FISH on an inversion candidate.** Two probes (p102 and p108) were designed on the potential inversion and labeled with green and red fluroscent colors. The CentC probe was used to located the centromere. The result indicated that both A188 and B73 had the same order of p102 and p108, which did not support the inversion.

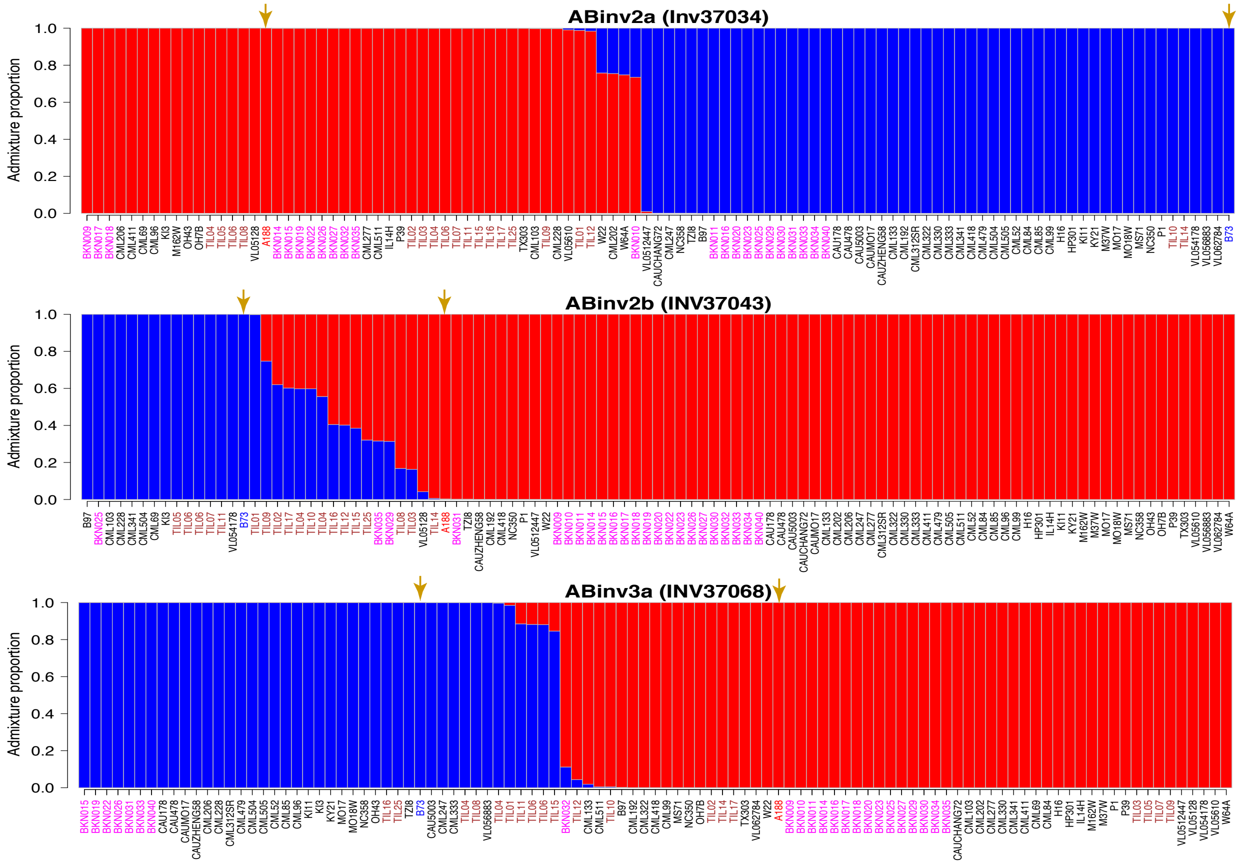


**Supplementary Figure 19. Structure analysis of three inversions in the maize Hapmap2 population.** The x-axis represents the maize lines. The y-axis represents the admixture proportion of two sub-populations for each line. Arrows point at A188 and B73. The maize wild ancesters, teosinte lines, are highlighted in brown, and landrace lines are highlighted in magenta.

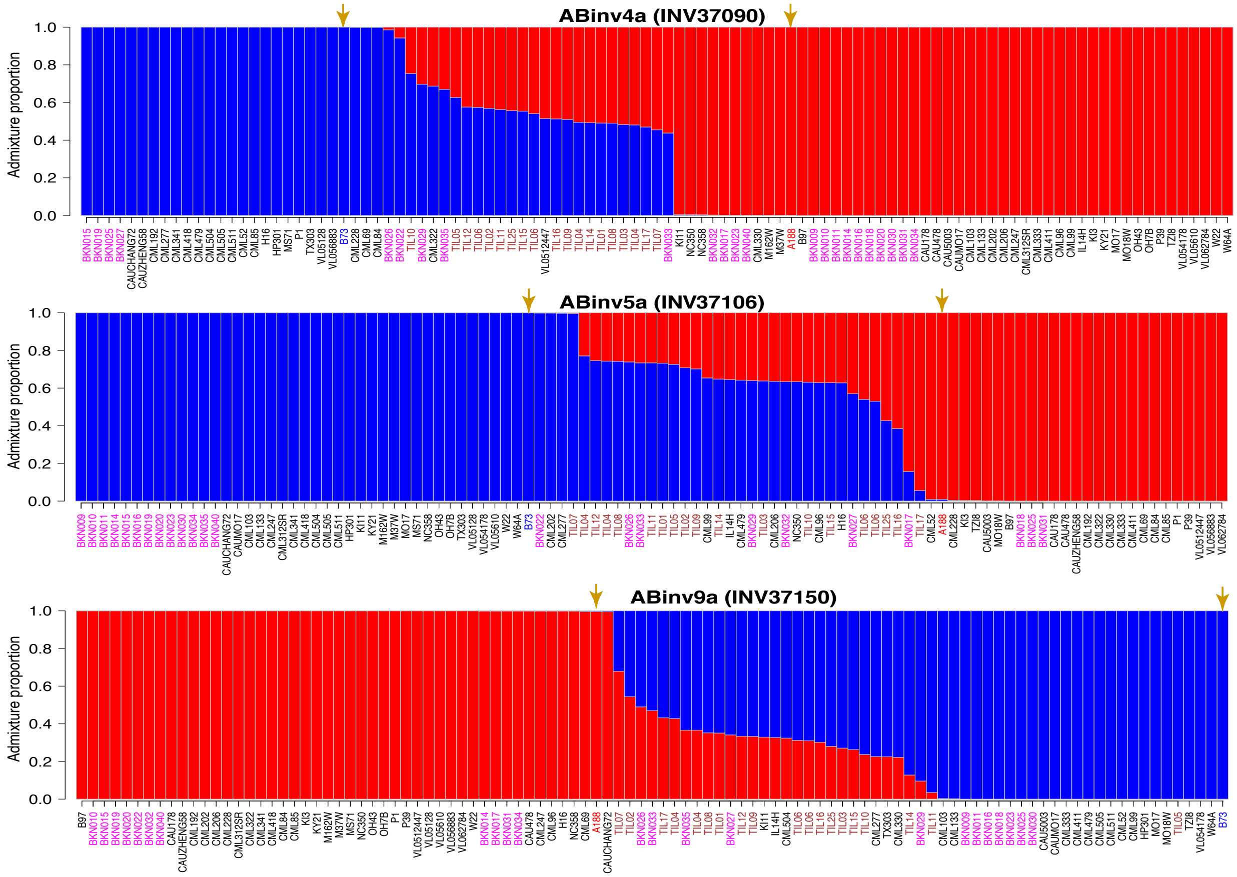


**Supplementary Figure 20. Structure analysis of other three inversions in the maize Hapmap2 population - II.** The x-axis represents the maize lines. The y-axis represents the admixture proportion of two sub-populations for each line. Arrows point at A188 and B73. The maize wild ancesters, teosinte lines, are highlighted in brown, and landrace lines are highlighted in magenta.

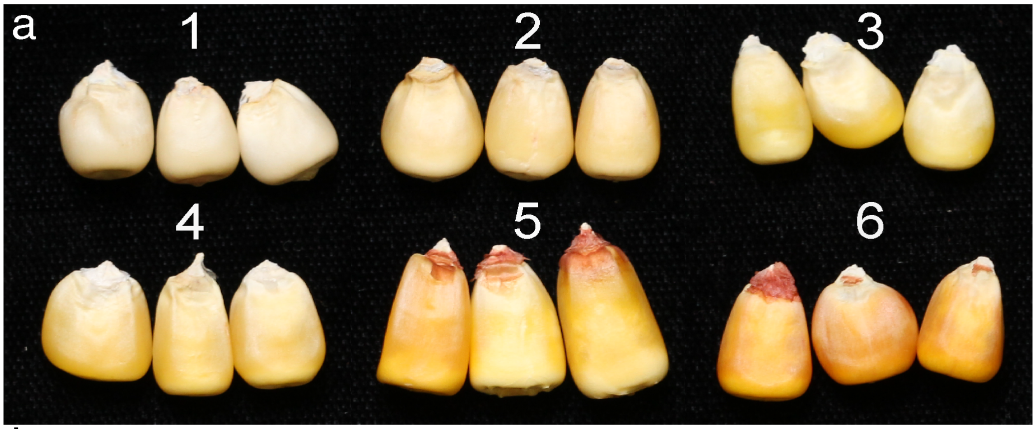


b

| Genotype combination of DHs | | | DH count of color codes from 1-6 | | | | | |
|---|---|---|---|---|---|---|---|---|
| **QTL_chr2** | **QTL_chr6** | **QTL_chr9** | **1** | **2** | **3** | **4** | **5** | **6** |
| A | A | A | 13 | 1 | 0 | 0 | 0 | 0 |
| B | A | A | 7 | 6 | 5 | 0 | 0 | 0 |
| A | A | B | 3 | 0 | 4 | 0 | 0 | 0 |
| B | A | B | 5 | 1 | 5 | 0 | 1 | 1 |
| A | B | A | 5 | 4 | 4 | 0 | 1 | 0 |
| B | B | A | 0 | 3 | 11 | 6 | 2 | 1 |
| A | B | B | 0 | 0 | 1 | 0 | 4 | 0 |
| B | B | B | 0 | 2 | 1 | 1 | 13 | 8 |

**Supplementary Figure 21. Phenotypic and genotypic data of DH lines. a**) A standard of kernel colors coded from 1-6. **b**) Counts of DH lines with kernel colors matching to standard codes for each genotype combination of three kernel color QTLs.

```
A188 1   MAIILVRAASPGLSAADSISHQGTLQCSTLLKTKRPAARRWMPCSLLGLHPWEAGRPSPA 60
B73  1   MAIILVRAASPGLSAADSISHQGTLQCSTLLKTKRPAARRWMPCSLLGLHPWEAGRPSPA 60

A188 61  VYSSLAVNPAGEAVVSSEQKVYDVVLKQAALLKRQLRTPVLDARPQDMDMPRNGLKEAYD 120
B73  61  VYSSLAVNPAGEAVVSSEQKVYDVVLKQAALLKRQLRTPVLDARPQDMDMPRNGLKEAYD 120

A188 121 RCGEICEEYAKTFYLGTMLMTEERRRAIWAIYVWCRRTDELVDGPNANYITPTALDRWEK 180
B73  121 RCGEICEEYAKTFYLGTMLMTEERRRAIWAIYVWCRRTDELVDGPNANYITPTALDRWEK 180

A188 181 RLEDLFTGRPYDMLDAALSDTISRFPIDIQPFRDMIEGMRSDLRKTRYNNFDELYMYCYY 240
B73  181 RLEDLFTGRPYDMLDAALSDTISRFPIDIQPFRDMIEGMRSDLRKTRYNNFDELYMYCYY 240

A188 241 VAGTVGLMSVPVMGIASESKATTESVYSAALALGIANQLTNILRDVGEDARRGRIYLPQD 300
B73  241 VAGTVGLMSVPVMGIATESKATTESVYSAALALGIANQLTNILRDVGEDARRGRIYLPQD 300

A188 301 ELAQAGLSDEDIFKGVVTNRWRNFMKRQIKRARMFFEEAERGVTELSQASRWPVWASLLL 360
B73  301 ELAQAGLSDEDIFKGVVTNRWRNFMKRQIKRARMFFEEAERGVTELSQASRWPVWASLLL 360

A188 361 YRQILDEIEANDYNNFTKRAYVGKGKKLLALPVAYGKSLLLPCSLRNGQT 410
B73  361 YRQILDEIEANDYNNFTKRAYVGKGKKLLALPVAYGKSLLLPCSLRNGQT 410
```

**Supplementary Figure 22. Alignment between Y1 protein sequences of A188 and B73.** Protein sequences of the transcript Zm00056a032392*T003 (A188) and the transcript Zm00001d036345*T001 (B73) were compared. Of 410 amino acids, 409 were identical. The polymorphic site is highlighted in red.

```
A188y1 TTCCAGGTGTCACCACTCAGCGTCCTCCGAACACAGGAGAGTCATGCGAT
B73y1  TTCCAGGTGTCACCACTCAGCGTCCTCCGAACACAGGAGAGTCATGCGAT

A188y1 GCGAGCTTGGCGATAAGCTTATCTATCCGCACCGCGTCTTCCTTCCTCCT
B73y1  GCGAGCTTGGCGATAAGCTTATCTATCCGCACCGCGTCTTCCTTCCTCCT

A188y1 GGGCGACCGGCCCTTCTTCTCTCCACGTCTCTCCCCCCTTCTTTCTCCAG
B73y1  GGGCGACCGGCCCTTCTTCTCTCCACGTCTCTCCCCCCTTCTTTCTCCAG

A188y1 AGCGAGCGTACGTATGCTACACACAGCAACAGCACAACAGTACTAGTTCC
B73y1  ACCGAGCGTACGTATGCTACACACAGCAACAGCACAACAGTACTAGTTCC

A188y1 ACCACAAGAAGATGCCCAATGCAAAGAAATAACCCATGCTTCTTGTCGAC
B73y1  ACCACAAGAAGATGCCCAATGCCAAGAAATAACCCATGCTTCTTGTCGAC

A188y1 GATCCAGCCGCAC-------------------------------------
B73y1  GATCCAGCCGCACTAGAGATGGCCAAACGGGCCGGCCCGGCCCGGCCCGG

A188y1 --------------------------------------------------
B73y1  CCCGGGCCCGGTGAAGCCCGGCCAAAACCGGGCCGGGCCTGCTGAGCCAG

A188y1 --------------------------------------------------
B73y1  CGGGCTTAAGTTTCTGTCCAAGCCCGGCCCGCAGCGGGCCTAAACAGGCC

A188y1 --------------------------------------------------
B73y1  GGGCCGGCCCGTTTAGCACGAAAAAACGGGCCAAAAAGCGGGCTAAACGG

A188y1 --------------------------------------------------
B73y1  GCCGGTAAGCACGTTTTAGTGTAAAAAAAACGGGCTTAACGGGCTTAGAG

A188y1 --------------------------------------------------
B73y1  GTAAACGGGCCGTGCCGGGCTAGCCCGCCGTGCCTAGTTTCCTGTCCAAG

A188y1 --------------------------------------------------
B73y1  CCCGCCCGCTTATTCTACCGTGCCGGGCTCGGACCGGGCCCAAAAAGCGG

A188y1 --------------------------------------------------
B73y1  GCTTCGTGCCGGGCTCACGGGCCTCGTGCTTTTTGGCCATCTATGAGCCG

A188y1 ---ACTTAGCATACGTACGCAAGAAGAGGAGAGGCCGGAGGTGCGCGTGC
B73y1  CACACTTAGCATACATACGCAAGAAGAGGAGAGGCCGGAGGTGCGCGTGC

A188y1 TCCTTGCTGTTCTGCTGACTGGTCTCATCATCTCATCCCACCACCACCAT
B73y1  TCCTTGCTGTTCTGCTGACTGGTCTCACCATCTCATCCCACCACCACCAC

A188y1 CACCA------TCTCTAGGATAAGATAGCAAATATATGGCCATCATACTC
B73y1  CACCACCACCATCTTTAGGATAAGATAGCAAATATATGGCCATCATACTC
[ … ]

A188y1 AATCGCTACTGCTCCCATGTTCATTGAGAAATGGCCAGACCTAGCCACCA
B73y1  AATCGCTACTGCTCCCATGTTCATTGAGAAATGGCCAGACCTAGCCACCA

A188y1 GAGAAGCTGCAATGCAAGGTTCAGGTTAGGCTAGATAGAAAGTTAAATGG
B73y1  GAGAAGCTGCAATGCAAGGTTCAGGTTAGGCTAGATAGAAAGTTAAATGG

A188y1 GGCAACATCACGAGGCCTTGATGAAAAACAGACAACCTGGTGAATTGTTG
B73y1  GGCAACATCAGGAGGCCTTGATGAAAAACAGACAACCTGGTGAATTGTTG

A188y1 TTGGGGTCAGGCACAGAACAGATAAGAGCCGCGCAGCCAACCTAGGGCTT
B73y1  TTGGGATCAGGCACAGAACAGATAAGAGCCGCGCAGCCAACCTAGGGCAT

A188y1 GTTCGGTT------AGCTCT------------------------------
B73y1  GTTTGGTTTCAATTAGTTCTAGGACTAAACTTTAGTCCTAGGACTAAACT

A188y1 -CAATCCATGTGGATTGAGT----GGGATTGTATGGGTTTGAAACCCA--
B73y1  TTAGTCCCTATATGTTTGGTTCTAGGGACTAAATAGATTCTAAAGTCATT

A188y1 -------------AACAAGTCAAACCTCTT---------CTCAT------
B73y1  AAATACATTGTCCAAAGACTCAAATACCCTTAGAATATACTCATGATATT

A188y1 --------------------------------------------------
B73y1  AGTTATCTATAAAAAGGTAAGGGCAACATGATAATTATGAGCTTTTAGTC

A188y1 TTTTTT------------------------CCAATCCCATCCAATCCAT-
B73y1  TCTTTTAGCACCTATGTGAAGGACTAAAGACTAAATCATTTTAGTCCATA

A188y1 -----------GTGTTTCGGAA----------------------------
B73y1  TTTTAGTCCTAGTGTTTGGCAAAAAAGGGACTAAAAGGGACTAAAAACTA

A188y1 ---------------------TAACCGAACAAGCCCCTAGATGGATACGG
B73y1  GAGACTAATCTTTAGTCCCTCTAACCAAACACCCCCCTAGATGGATACGG

A188y1 AACATTCGCCTCTTATTCGGAGCAATATATGTCTCTCAAGGAAAGAGCCC
B73y1  AACATTCGCCTCTTATTCGGAGCAATATATGTCTCTCAAGGAAAGAGCCC

A188y1 AACATGTATACTGCCTTCTTTTTCTCATCCCAGATTTGGGGGAAAAACAA
B73y1  AACATGTATACTGCCTTCTTTTTCTCATCCCAGATTTGGGGGAAAAACAA

A188y1 TGTAAATGCCAATGGTATCGTAGGAAGATTACTAGAAGTAAATGCCAATG
B73y1  TGTAAATGCCAATGGTATCGTAGGAAGATTACTAGAAGTAAATGCCAATG

A188y1 TAAAAACAGATGAGTTGGCATTTACATGATAGGATGGTGGGATCATCAGA
B73y1  TAAAAACAGATGAGTTGGCATTTACATGATAGGATGGTGGGATCATCAGA

A188y1 CTGAAAATGATAGGGGATTGTGCTCCCCTGCGACTCCAACTACTAAACAA
B73y1  CTGAAAATGATAGGGGATTGTGCTCCCCTGCGACTCCAACTATTAAACAA
```

**Supplementary Figure 23. Alignment of 5' and 3' flanking sequences of A188 *y1* and B73 *Y1* alleles.** Translation start sites and translation termination sites are highlighted in yellow. A (CCA)n microsatellite variation at the 5' untranslated region is highlighted in green. Most gene body sequences are skipped.

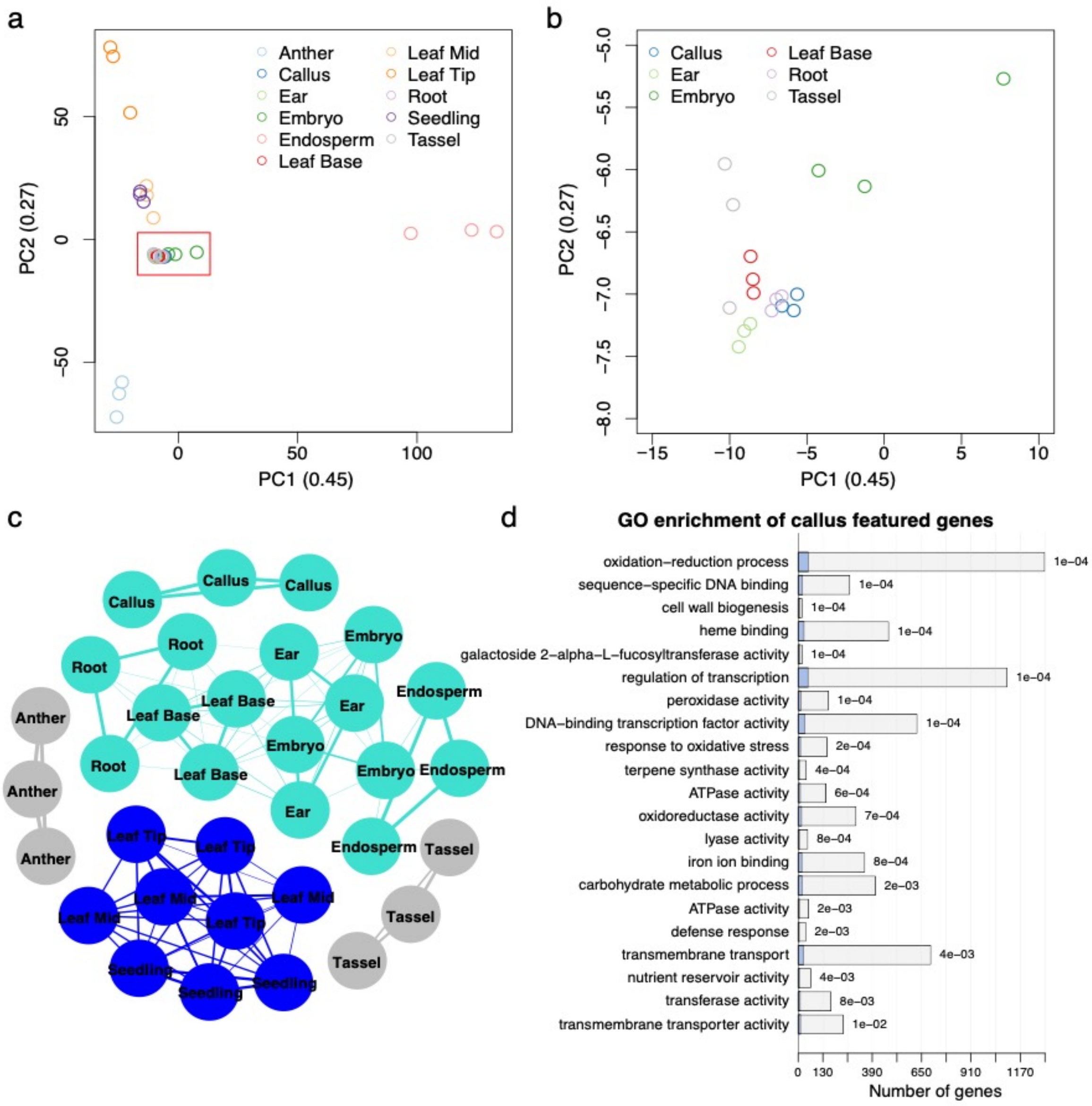


**Supplementary Figure 24. sample clustering and callus-featured genes a**) Pincipal component analysis (PCA) results of gene expressions in 11 A188 tissues. The x-axis and y-axis represent the first component (PC1) and the second component (PC2), respectively. The numbers within the parentheses stand for the proportions of the variation of gene expressions explained by either PC1 or PC2. **b**) Enlarged PCA plot of the red box in **a**. **c**) The network of 33 RNA-Seq samples from 11 tissue types constructed based on their gene expression. Two major clusters were identified. One cluster includes callus, root, leaf base, ear, embryo, and endospeam; the other cluster includes leaf tip, leaf middle, and seedling. The leaf base, middle, and tip are three parts from base to tip from the same leaf. **d**) GO enrichment of callus featured genes. In each barplot, a blue bar stands for the number callus featured genes and the whole bar (blue and empty) stands for the total number of genes of the associated GO term. P-values were labeled on the top of each bar. Only the GO terms with the p-value smaller than 0.01 and containing at least five callus featured genes were plotted.

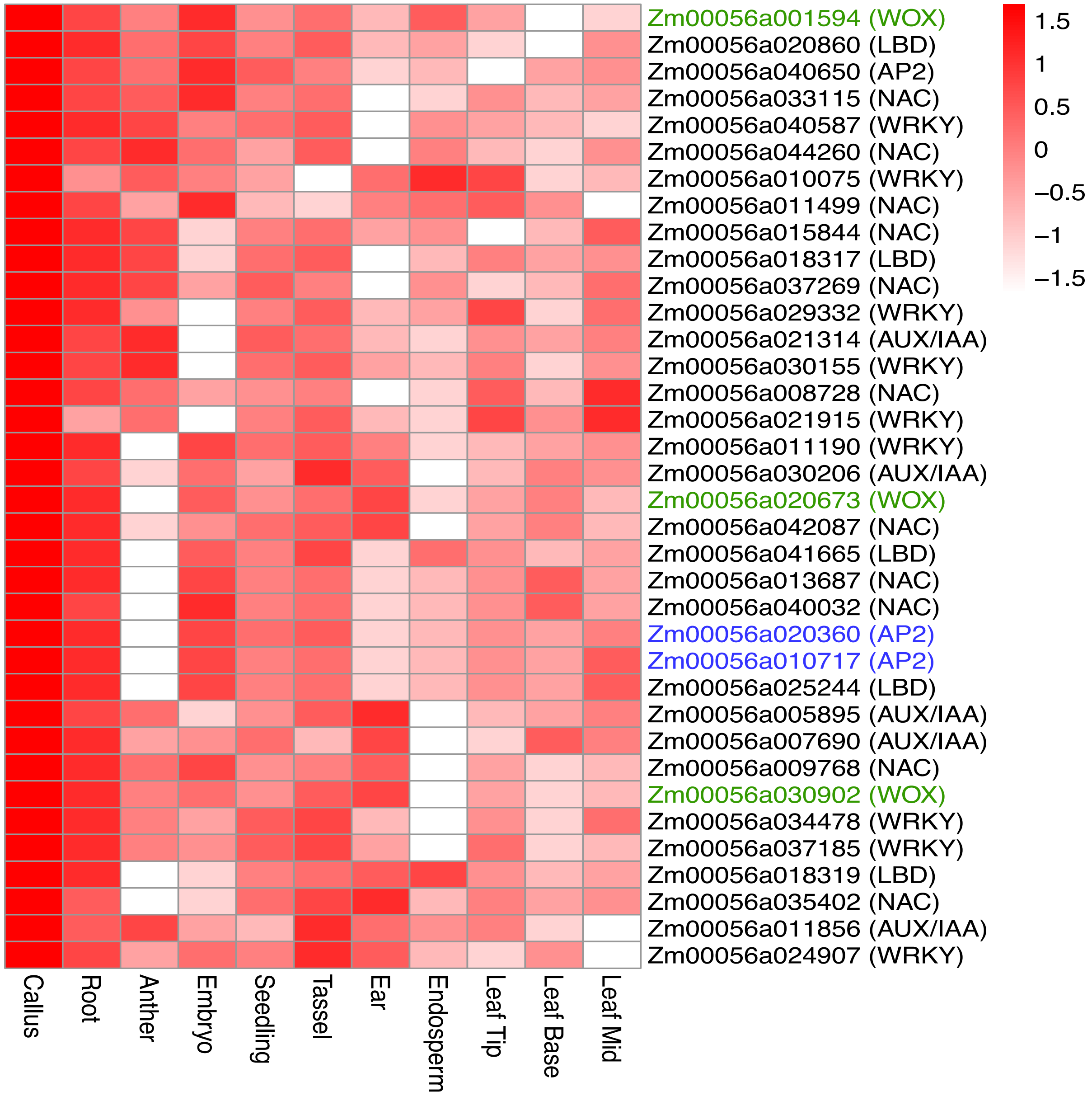


**Supplementary Figure 25. Heatmap plots of expression of callus-featured TF genes.** The row and column represent the callus-featured TFs and tissues, respectively. The values of gene-wise quantile-quantile normalized (qqnorm) gene expressions are color-coded. The qqnorm implemented by an R package “qqnorm” normalized gene expressions to a Gaussian distribution. Genes homologous to *Baby boom* and *Wuschel2* are colored in blue and green, respectively.
